# The “Grove-First” Framework: Starting in the Grove to Find Therapies for Huanglongbing

**DOI:** 10.1101/2025.05.22.655577

**Authors:** Michelle L. Heck, Nicholas R. Larson, Guilherme Locatelli, Ellen F. Cochrane, Samuel Coradetti, Lukas Hallman, Lynn Johnson, W. Cody Estes, Ariana Makar, Nursena Demirden, Marco Pitino, Robert G. Shatters, Robert C. Adair, Frank Giles, John-Paul Fox, Douglas Stuehler, Joanne Hodge, James Hoffman, Victoria Blake, Loushendy Ulysse, Lisvet Ramirez-Barrera, Riley McKenna, Luke Thompson, Lucy Bennett, Adriana Larrea-Sarmiento, Alejandro Olmedo-Velarde, Samantha Barkee, Chase Weeks-Purdy, Flavia Zambon, Ketan Shende, Lorenzo Rossi, Tom D’Elia, Mark A. Ritenour, Brian T. Scully, Randall P. Niedz

## Abstract

Citrus greening disease, also known as huanglongbing (HLB), is the most serious vector-borne bacterial disease of citrus world-wide^1,2^. There is an immediate global need to provide the citrus industry with relief from HLB and a return to profitable citrus production. Standard screening methods for HLB therapeutic treatments typically involve various laboratory-based assays to select treatments with antimicrobial properties^3^, which then advance to greenhouse and eventually field testing in a workflow that takes multiple years. Unlike traditional lab-first screening, we present a design of experiments framework^4,5^, referred to as Grove-First, that rapidly screens treatments with regulatory-friendly profiles in commercial citrus groves using trunk injection to select treatments that improve tree health and fruit yield over the course of a single growing season. Using this framework, we identified candidate treatments with effects comparable to or better than the standard oxytetracycline (OTC) on visual tree-health and/or yield indices in an initial screen of HLB-positive 8-year-old Valencia sweet orange trees. Expanded trials in commercial citrus groves allowed us to validate the initial screening results at other locations and in other citrus varieties. Grove-First rapidly accelerated the identification and large-scale field testing of HLB therapies, some of which are available for growers to use immediately and others that require further field testing and/or regulatory actions.

## Main

HLB is a fatal disease in citrus associated with the unculturable Gram-negative α-Proteobacteria: ‘*Candidatus* Liberibacter asiaticus’ (*C*Las), found in Asia, North and South America, Oceania and the Arabian Peninsula^1,6^; ‘*C*. Liberibacter americanus’ (*C*Lam), in South America^7,8^; and ‘*C*. Liberibacter africanus (*C*Laf) found in Africa and the Arabian Peninsula^9,10^. The bacteria are restricted to the plant phloem vascular tissue, and their natural spread is primarily by two species of psyllids, hemipteran insect vectors^2,11,12^. While different species of citrus and citrus varieties vary in HLB symptomology, infected trees decline over time. Symptoms typically manifest as yellowing of leaves, tufting, decline of root biomass, asymmetric blotchy mottling, reduction in juice °Brix (leading to a bitter taste), uneven and incomplete fruit ripening, thinning of the canopy, premature fruit drop and ultimately tree death^12^. Hence, HLB negatively affects citrus production worldwide. In some areas, such as Florida in the southern United States, the disease has decimated the industry since its detection in 2005^13^. Trunk injection of oxytetracycline (OTC) became a widespread practice among Florida growers in 2023 after two injectable OTC products, ReMedium TI^TM^ and Rectify^TM^, received a Federal Insecticide, Fungicide, and Rodenticide Act (FIFRA) Section 24(c) Florida special local need registration from the Environmental Protection Agency (EPA) in October 2022 for HLB management^14–18^. The swift and efficient adoption of trunk injection by citrus growers in Florida challenges the notion that treating individual trees is economically infeasible^19^.

Treatments that show promise in laboratory or greenhouse assays often fail to deliver consistent or useful improvements when tested in commercial citrus groves^20–22^ because laboratory results do not reliably predict field performance under the complex and variable conditions encountered in citrus production^21,23^, treatment effects are not sufficiently large enough to have an economic impact^22,24^ and other reasons. These disconnects may also occur because lab assays often focus narrowly on antimicrobial activity or surrogate outcomes^3,25^, while field efficacy depends on a broader range of factors-including systemic movement, environmental interactions, and the ultimate impact on fruit yield and quality-that are not captured in controlled experiments. Moreover, the cost associated with advancing synthetic pesticides to market is substantial^26^.

Finding, testing, and validating potential solutions for citrus HLB is inherently challenging due to the crop’s multi-year growth cycle, perennial nature, and the variability of environmental conditions and management practices. Newly planted trees require three to four years to produce fruit and eight years to reach full productivity, and established groves have one crop annually. Because the sole metric of success is the consistent and profitable production of high-quality fruit, collecting the data required to validate a solution is inherently a multi-year process. This type of system results in a dilemma for the growers: adopting untested or inadequately tested solutions could lead to costly failures^20,27^, while waiting for definitive proof risks losing their groves to HLB. Grove-First was developed specifically to address this dilemma by enabling researchers and growers to test potential solutions directly in commercial groves, under real-world conditions and across multiple seasons and locations. By prioritizing field-based evaluation of fruit yield and quality, the only metrics that define economic viability, Grove-First (**Fig. 1a**) bridges the gap between laboratory research and practical application. This collaborative approach accelerates the identification of effective, scalable therapies by focusing on what works in the grove and empowers the citrus industry to respond more rapidly and with higher confidence to the urgent need to solve HLB.

**Fig. 1.**
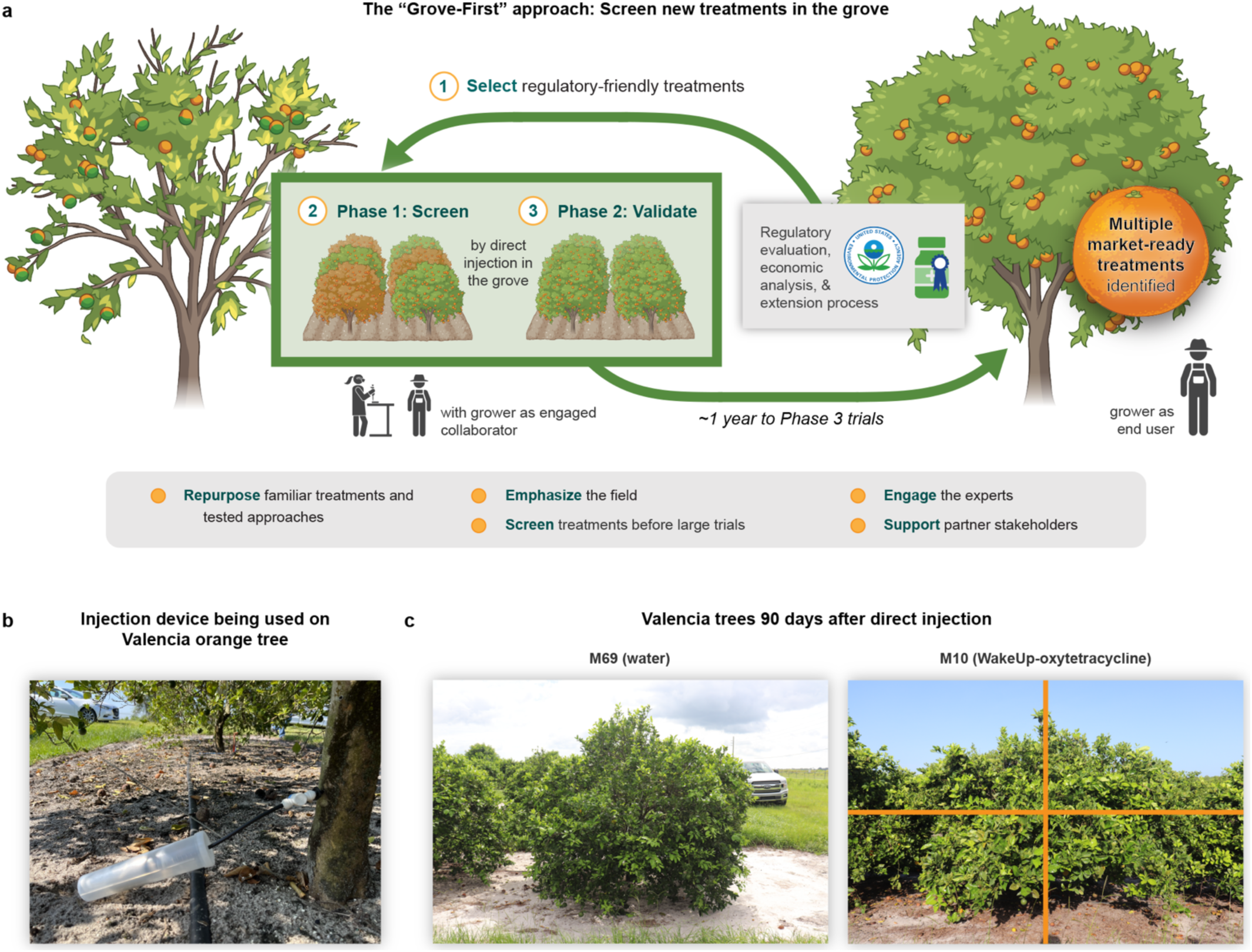
The “Grove-First” approach: Screening new HLB therapies in the grove. (a) Conceptual overview of the Grove-First framework used to rapidly identify treatments to rejuvenate trees affected by huanglongbing (HLB). Regulatory-friendly candidate treatments are screened directly in the grove (Phase 1) and validated (Phase 2) through direct trunk injection into HLB-affected citrus trees. This approach prioritizes field efficacy, rapid screening, and active grower collaboration to accelerate the development of commercially viable solutions. (b) FlexInject^TM^ injector (TJ Biotech) used for injection delivery of candidate treatments into citrus trees. (c) Visual comparison of HLB-affected Valencia sweet orange trees (*Citrus sinensis* L. Osbeck) 90 days after injection. Left: untreated control tree, right: tree treated with M10 (combination of WakeUp and Rectify^TM^ oxytetracycline at 1% and 10,000 ppm, respectively).

The Grove-First strategy is a multi-phase, collaborative approach to rigorously screen treatments, ensuring that only treatments with large, unmistakable effects advance to broader citrus industry use. While the Grove-First strategy can apply to any type of treatment, this report describes its application to screen chemistries to manage HLB using direct trunk injection (**Fig. 1b**) and evaluating treatment effects on tree health (**Fig. 1c**), fruit yield (**Fig. 2**), and fruit quality (**Fig. 3**). The focus is on effects that are large and unmistakable, which are necessary to grow citrus profitably under HLB pressure. The use of trunk injection was inspired by Florida citrus growers, who demonstrated that the approach was economically feasible at commercial scale during the 2023 and 2024 growing season.

**Fig. 2.**
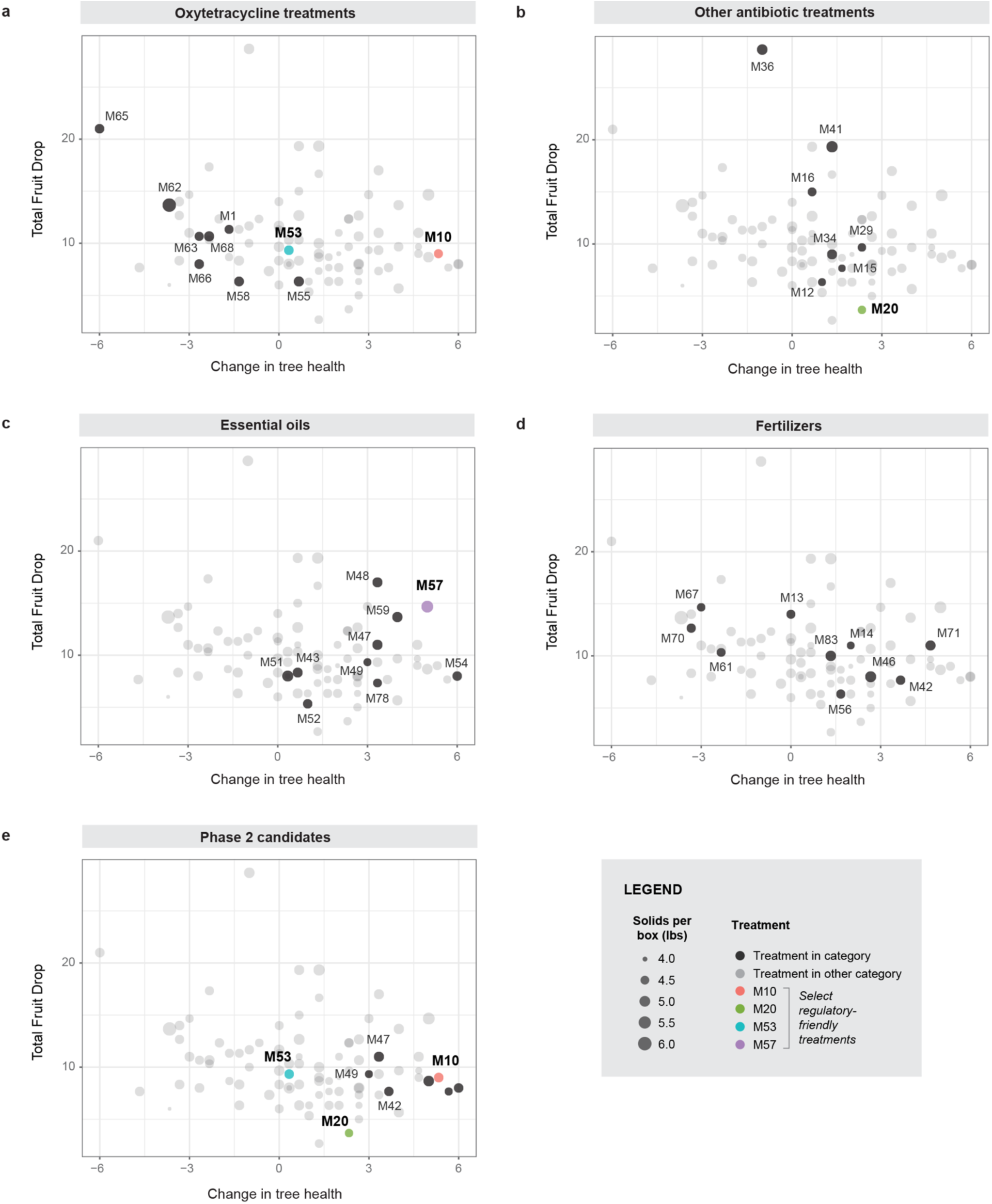
Tree-health improvement, fruit drop, and solids content from the Phase 1 screen for candidate treatments. Bubble-plot panels compare the 180-day change in tree-health index (x-axis) with the total number of fruit dropped per tree (y-axis) for every treatment screened in Phase 1 (one bubble = mean of three trees). Bubble cross-sectional area is proportional to pounds of soluble solids per standard field box, providing a third yield metric. Each panel highlights a different treatment class in the context of the performance of all treatments: (a) readily deployable oxytetracycline formulations, (b) additional antibiotics, (c) essential oils, (d) fertilizers, and (e) the treatments advanced to Phase 2.

**Fig. 3.**
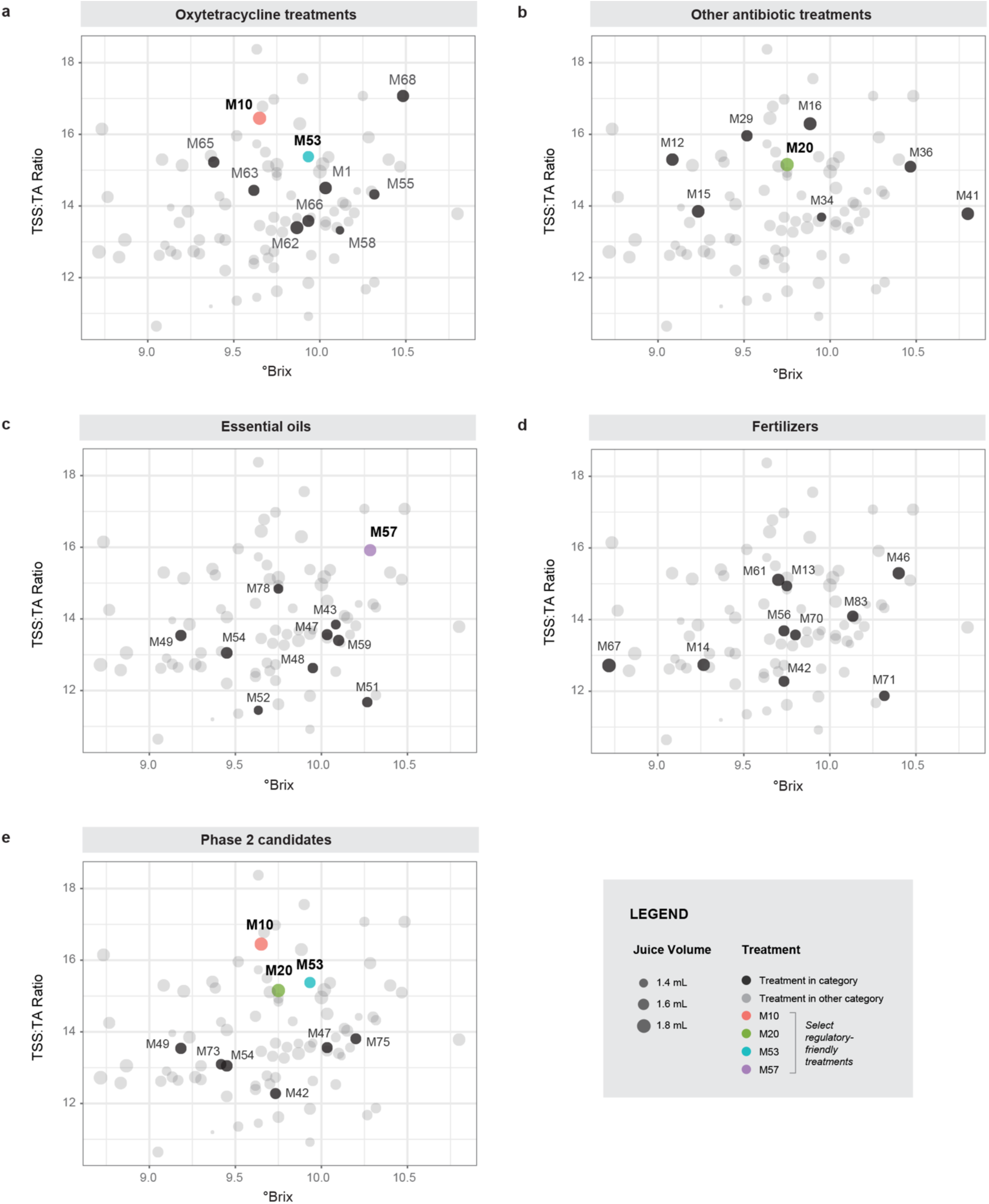
Internal fruit quality analysis for candidate treatments from the Phase 1 screen. Panels (a-e) plot °Brix (x-axis) and the total-soluble-solids-to-titratable-acidity ratio (TSS:TA; y-axis) for the identical treatment means shown in Fig. 2. Bubble size scales with corrected juice volume (mL/fruit), visualizing sweetness/acidity to juice yield for selected treatments in the context of the performance for all treatments. Color coding matches Fig. 2, enabling direct visual comparison between external yield metrics (Fig. 2) and internal quality traits (Fig. 3) across (a) oxytetracycline variants and combinations, (b) other antibiotics, (c) essential oils, (d) fertilizers, and (e) the Phase 2 treatment subset.

The Grove-First strategy enables screening a large number of treatments to control HLB using a three-phase approach designed to efficiently identify effective treatments. Phase 1 initiates with a small-scale screening trial, where each treatment is injected into *n* = 3 selected trees in a single grove to identify those that have large effects on the trees (**Fig. 1c**). This initial phase ensures that only promising chemistries progress to Phase 2, thereby saving resources by eliminating ineffective treatments early on. Phase 2 involves grower validation trials conducted in multiple commercial groves, where multiple trees within a row (or multiple rows, if feasible) are treated for grower assessment of efficacy. Treatments that show large effects in Phase 2 validate the signal observed in Phase 1 and proceed to Phase 3, which includes large-scale production testing, assessment of treatment robustness under varying conditions, and grower adoption. This phased approach tolerates an initial high false positive rate (type 1 error) to avoid discarding promising treatments with large effects in Phase 1, while narrowing down treatments through the successive phased trials to ensure that resources are efficiently allocated towards the most effective treatments. The focus on large effects is particularly useful given the severity of HLB where small incremental improvements are unlikely to restore economic viability to HLB-affected citrus groves. There is an urgent need for solutions to HLB in the citrus industry. Growers need a broader array of treatments to further improve economic yield and hedge against the emergence of OTC-resistant *C*Las or future regulatory restriction of OTC use for HLB management.

Phase 1 screening for large effects prioritizes therapies that have the potential to produce significant, easily observable improvements in tree and fruit health. This strategy is analogous to field evaluation of germplasm for phenotypes of interest in a standard plant breeding program. Similarly, in the context of a treatment for HLB, even a single tree showing a remarkable positive response may justify the resources to advance that treatment to further trials to validate the effect^28^. A consequence of screening for large effect sizes is that the approach is extremely efficient and does not require a large sample size for the initial Phase 1 screening trial. We hypothesized that a sample size of three is sufficient for this first phase screening per the concept of statistical power where a large effect size compensates for a small sample size^29^. The need for such a large effect illustrates the seriousness of this disease that has decimated Florida’s citrus industry and why only treatments with very large and robust effects are commercially viable.

The increasingly stringent evaluation phases use tree health and fruit metrics that align with citrus grower assessments and with Florida’s official citrus quality standards as defined by state regulations (§ 601.11, Fla. Stat. (2024)). By orienting towards a plant breeding-type approach focused on validating therapies in citrus groves managed by citrus growers, Grove-First specifically accounts for the challenges that citrus farmers routinely face - the urgency to find a treatment, financial constraints, and U.S. regulatory laws affecting HLB control measures. Our study shows that Grove-First is a systems-level approach, involving scientists, growers, citrus grower organizations, regulatory agencies, state and federal funding agencies, agribusinesses, technology transfer specialists, and even high school students ensures the rapid evaluation and delivery of practical, impactful solutions to HLB management.

### Treatment impact on tree health over the growing season

Treatments were selected across a range of chemistries from a therapy repurposing list generated internally or submitted to our team under a confidentiality agreement from other researchers and growers interested in participating in the screen (**Extended Data Table 1**). Each treatment is given an “M” identification number (**Extended Data Table 1**). In Phase 1, we screened 88 treatments (**Extended Data Table 1**) by injection into 8-year-old Valencia sweet orange trees at the USDA Agricultural Research Service Picos Research Farm in Fort Pierce, FL, 67 of which had no antimicrobial activity in a plate-based assay testing growth inhibition of *Agrobacterium tumefaciens* (**Extended Data Table 1**). We used three trees per treatment in trees that were randomized within a block containing eight rows of trees. To minimize biological variation within treatments, trees were classified as alpha, beta or omega. Alpha trees displayed minimal HLB disease symptoms and largest trunk diameters, whereas omega trees had more severe HLB symptoms and smaller trunk diameters (**Extended Data Table 1, Extended Data Fig. 1a**). While there was no obvious bias in starting tree health ratings across rows 1-6, rows 7 and 8 had no alpha-ranked trees and more omega-ranked trees relative to the other rows (**Extended Data Fig. 1a**). A Pearson’s χ² test of independence with Monte Carlo resampling (9,999 replicates) showed that the distribution of tree ratings differed significantly across rows (χ² = 55.53, simulated *P* = 0.0001). Each treatment was only delivered into one class of tree and the three trees per treatment were randomized throughout the block to minimize row biases on tree response. The Rectify^TM^ label formulation of OTC (AgroSource, Inc.) was injected as a positive control (M53) against which all phenotypic changes were measured in the Phase 1 trial. Trunk damage at the sites of injection ranged from no or mild damage (**Extended Data Fig. 2a**), moderate damage (**Extended Data Fig. 2b**) to severe damage which led to splitting of the trunk (**Extended Data Fig. 2c**). Most injection sites sustained mild or no damage (**Extended Data Fig. 2a**). We did not identify any patterns that would predict the development of trunk damage.

Phenotypic effects after injection were observed quickly for some treatments. Within seven days post-injection, some treatments displayed large phytotoxic effects (**Extended Data Fig. 3a,b**). For a subset of those treatments, the phytotoxicity resulted in leaf drop followed by the re-growth of young citrus tissue (**Extended Data Fig. 3b**). These unexpected observations were an early indicator that treatment-specific, tree-level responses would occur. Thus, we recorded changes in tree health index at regular intervals (90-, 180- and 270- days, **Extended Data Fig. 4**). At day 90-post injection (**Extended Data Fig. 4a**), 10 treatments had the same change in tree health index as OTC at pH 2 (M53) and 22 treatments showed larger, positive effects on tree foliar health as compared to OTC at pH2 (M53). Two of the treatments showing qualitatively larger effects were fertilizer treatments, zinc nitrate hexahydrate (M71) and ammonium thiosulfate (M46), the latter showing the largest positive change in tree health (**Extended Data** Fig. 4a). Dipotassium EDTA (M73), a common chelating agent, and muriatic acid (M75), which is used to lower the pH of the injectable OTC products showed positive effects at 90-days post-injection. Two antibiotics, trimethoprim/sulfamethoxazole (SMZ-TMP, M29) and metronidazole (M34) had positive effects at day 90 post-injection. Notably, five of the ten of the essential oils we tested showed positive effects on tree health (**Fig. 2c**): star anise oil (M54), clove oil (M59), tea tree oil (M57), cinnamon oil (M52) and pongamia oil (M49). At day 180 post-injection, six treatments improved tree health similarly to OTC at pH2 (M53) and 21 treatments showed a greater improvement in tree health as compared to OTC at pH2 (M53). At 180-days post injection (**Extended Data Fig. 4b**), EDTA (M73) and muriatic acid (M75) treatments continued to show improvements in tree health. Additionally, six of the ten essential oils: star anise (M54), citronella (M48), clove oil (M59), pongamia oil (M46), camphor (M78), tea tree oil (M57); four fertilizers: zinc nitrate hexahydrate (M71), ammonium thiosulfate (M46), potassium phosphate monobasic (M42) and potassium permanganate (M14) and two antibiotics: OTC combined with the adjuvant WakeUp Advantage (M10) and cephalexin (M74) showed improvement in tree health. Day 270-post injection was during the early spring, when *C*Las titers would have been higher due to several months at cooler temperatures and when tree condition has been reported to decline^21,22^. At this time point, most treatments did not show tree improvement or tree condition even worsened (**Extended Data Fig. 4c**), including for OTC pH2 standard (M53). However, two antibiotic treatments: streptomycin in combination with OTC (M20) and OTC combined with WakeUp Advantage (M10), the fertilizer potassium phosphate (M42) and sodium benzoate (M40) continued to show improvement in tree health (**Extended Data Fig. 4c**). Among the 67 treatments that did not suppress bacterial growth in the plate assay (**Extended Data Table 1**), many of these showed a positive effect in tree health response over the growing season, including most of the essential oils (**Extended Data Fig. 4**). Overall, treatments delivered into omega-rated trees showed the largest positive change on tree health at 90- and 180-days post-injection (Welch’s ANOVA *P-*values < 0.0001, **Extended Data Fig. 5a,b**), which waned by 270-days post-injection (Welch’s ANOVA *P-*value = 0.1284; **Extended Data Fig. 5c**). Alpha-rated trees exhibited the smallest change over time, indicating a ceiling effect of our tree health rating system.

A clearly visible increase in leaf expansion was an early positive indicator that a treatment was positively improving tree health. The combination treatment of OTC and the surfactant WakeUp Advantage (M10) first showed visibly larger leaves by 90 days post-injection (**Fig. 1c, Extended Data Fig. 6a**), which persisted for the duration of the study; however, the large-leaf effect appeared sectored: leaf-size differences were significant in only the lower half of the tree on the east, injected side (two-way ANOVA east vs west × quadrant interaction, *P* = 0; Tukey post-hoc east Q3 > all west quadrants, *P* < 0.001, **Extended Data Fig. 6b**), suggesting that systemic distribution of the treatment throughout the tree was uneven.

Fruit drop is a problematic symptom of HLB, leading to premature abscission of immature or underdeveloped fruit and resulting in significant yield losses^12^. To evaluate the effectiveness of each treatment to mitigate fruit drop, we measured fruit drop over a 2-week period from each tree in the experimental plot. No treatments showed a statistically significant decrease in fruit drop as compared to OTC pH2 (M53, **Extended Data Table 2**), although the top performing treatments in terms of lowest median drop included the streptomycin and OTC combination (M20), potassium phosphate monobasic (M42) and D-leucine (M76, **Extended Data Fig. 7a**). As expected, trees initially rated omega tended to have the highest standard deviation across the fruit drop measures, independent of treatment (**Extended Data Fig. 7b**).

Looking across all the permutations of OTC formulations we tested for tree health (**Fig. 2a**) and fruit quality metrics (**Fig. 3a**), there were no consistent or predictable trends in tree performance based on dose rate. M68, which was OTC at pH2 at 5500 parts per million (ppm), or roughly half the label rate, showed higher °Brix as compared to M53, (10,000 ppm) for the tree size used in the experiment (**Fig. 3a**). OTC plus WakeUp (M10) had a higher ratio of total soluble solids to titratable acidity (TSS:TA ratio) as compared to M53 (**Fig. 3a**), suggesting that fruit on the M10 treated trees may have benefitted from a longer ripening time on the tree. Residue analysis of whole fruit revealed six out of 15 samples tested had a detectable level of OTC, with the highest detection level at 4 parts per billion (ppb, **Extended Data Table 1**), which is below the tolerance limit of 10 ppb in citrus. Of the other antibiotics tested, doxycycline (M36) and ciprofloxacin (M41), broad-spectrum antibiotics with activity against Gram-negative bacteria, showed impressive internal fruit quality metrics, high °Brix and TSS:TA ratio levels, whereas M20, the combination of streptomycin and OTC, showed a relatively high fruit weight as compared to the other treatments. All essential oils improved tree health at 180 days post-injection (**Fig. 2c**), with tea tree oil (M57), resulting in an impressive tree health, yield and fruit quality response on par with or better than some of the antibiotic treatments (**Figs. 2c, 3c and 4**). Among the fertilizers tested, zinc nitrate hexahydrate (M71), potassium phosphate monobasic (M42) and ammonium thiosulfate (M46) stood out in tree health, yield and fruit quality indices (**Fig. 2d, 3d**), with potassium phosphate monobasic (M42) exhibiting lower °Brix as compared to M71 and M46.

**Fig. 4.**
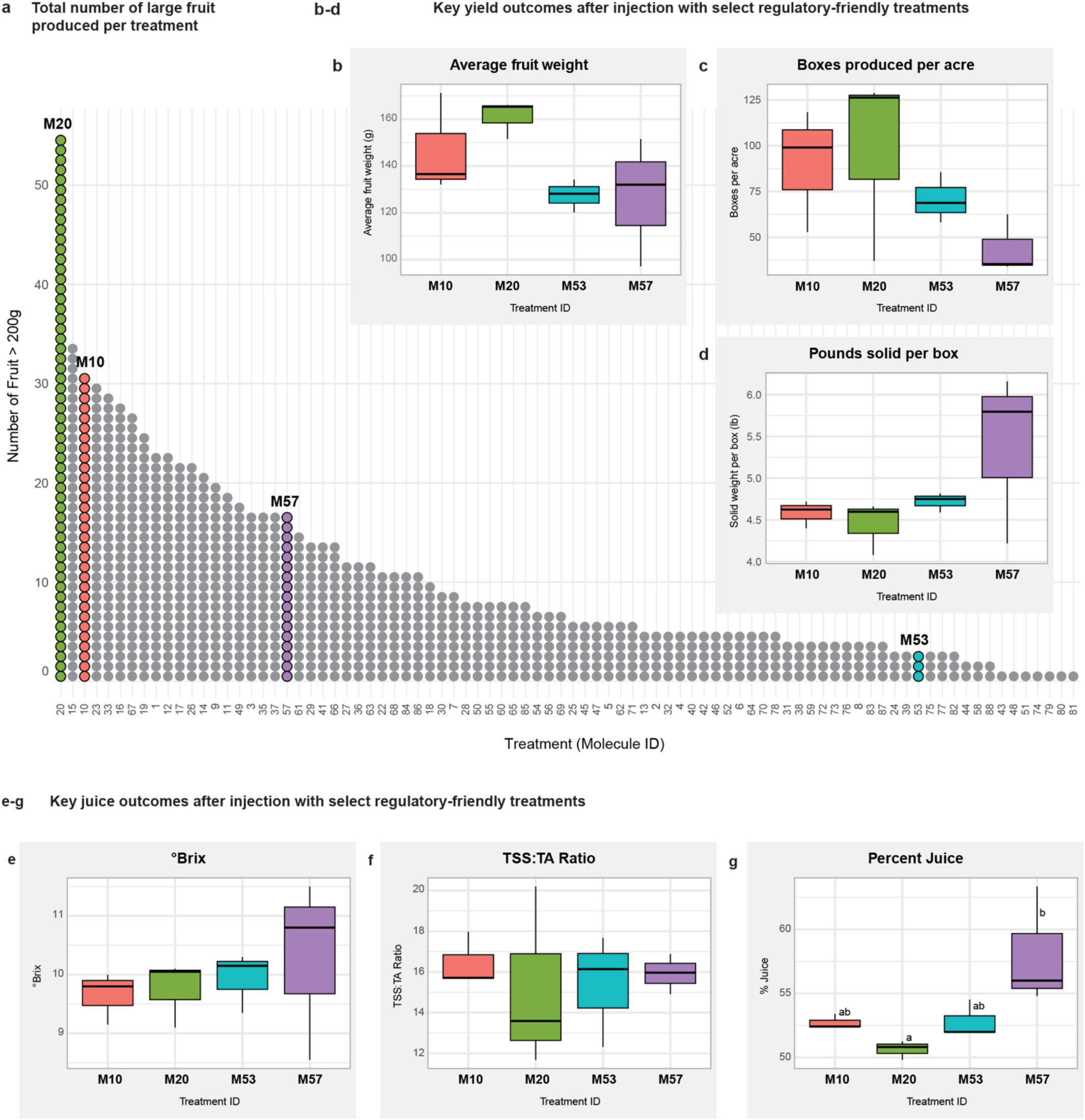
Large-fruit yield and detailed quality attributes of four leading trunk-injection treatments from the Phase 1 screen. (a) Pareto bar chart ranking every screened treatment by the number of fruit weighing > 200 g. Four treatments of interest, Rectify^TM^ oxytetracycline plus WakeUp Advantage (M10), Rectify^TM^ oxytetracycline control (M53), tea-tree oil (M57) and streptomycin plus Rectify^TM^ oxytetracycline (M20), are colored individually; all other treatments appear in grey. (b-g) Box-and-whisker comparisons of the same four treatments for six key performance metrics: (b) average fruit weight, (c) boxes per acre, (d) pounds solids per box, (e) °Brix, (f) TSS:TA ratio and (g) percent juice. Each box represents the mean response of the three replicate trees in the Phase 1 trial.

### Treatment impact on fruit yield and quality

Positive changes in tree health were not correlated with changes in the number of fruit (**Extended Data Fig. 8a**), but they were positively correlated with fruit weight (**Extended Data Fig. 8b**). A total of 54 treatments had a significant, positive effect on the average number of fruit per tree as compared to M53 (*P* <0.05, **Extended Data Fig. 9a, Extended Data Table 3**), but only two had a positive effect on fruit weight (*P*<0.05, **Extended Data Fig. 9b, Extended Data Table 4**, eight if you accept a significance level of alpha = 0.1). Of these, three were antibiotics: the OTC-streptomycin combination (M20), cefotaxime (M16) and OTC, pH2 with iron (M58). The row where the trees were planted had a significant impact on the number of fruit produced and fruit weight, independent of treatment (**Extended Data Fig. 1b**). A Kruskal–Wallis test revealed that the number of fruit per tree and fruit weight varied significantly among rows (*P* = 0, **Extended Data Fig. 1c**). There was a significant negative correlation between fruit weight and the number of fruit per tree (**Extended Data Fig. 9c**). No statistically significant increases were observed for °Brix, TSS:TA ratio or pounds solid per box measurements between any of the treatments and OTC pH2 (M53) tested in Phase 1 (**Extended Data Table 5, 6**). The tea tree oil treatment (M57) resulted in significantly higher average percent juice as compared to the OTC-streptomycin combination M20 (ANOVA *P* =0.02, **Fig. 4**). The streptomycin-OTC combination (M20), the WakeUp-OTC combination (M10) and the tea tree oil treatment (M57) all produced more fruit weighing over 200g (large fruit) as compared to OTC at pH2 (M53), (**Fig. 4a**). Only two treatments resulted in all three trees achieving °Brix levels over 10: doxycycline (M36) and ciprofloxacin (M41) (**Extended Data Fig. 10a**). These two antibiotics also showed low variance in their TSS:TA ratio, highlighting uniform internal fruit quality across all three trees receiving the treatments (**Extended Data Fig. 10b**).

While comparison of treatment effects to M53 (OTC) was the predominant approach we used to analyze the Phase 1 screening data, alternative approaches to analyze the data yielded similar insights. For example, hierarchical clustering analysis grouped the treatment effects into ten distinct clusters (**Fig. 5**), each comprising a unique subset of treatments with varying group sizes. While the clusters were not strictly defined by any single horticultural effect, two adjoining sub-clusters contained treatments that had the highest °Brix increase (**Fig. 5**), including tea tree oil (M57), ciprofloxacin (M41) and doxycycline (M36). The antibiotic treatments were predominantly distributed across most clusters, but the OTC-strep combination (M20), the OTC-WakeUp combination (M10), vancomycin (M15), and betadine (M15) clustered together. Treatments highlighted in bold (**Fig. 5**) represent those advanced to Phase 2 trials to validate the effects we observed in Phase 1. Another alternative to analyzing the data was by comparing treatment effects to the mean values for each tree class: alpha, beta and omega (**Extended Data Table 7**). Using this approach with a *P*-value cutoff of 0.1, a few treatment combinations stood out. For an increase in the °Brix values, these include the use of OTC at half the label rate (M68, 5500 ppm), ciprofloxacin (M41), 2% ethanol (M22), iron II nitrate nonahydrate (M83) and doxycycline (M36). Among other treatments, several essential oils had significant effects, including geranium oil (M47) on the number of fruit and pongamia (M49) and tea tree oil (M57) on the number of fruit > 200 g.

**Fig 5.**
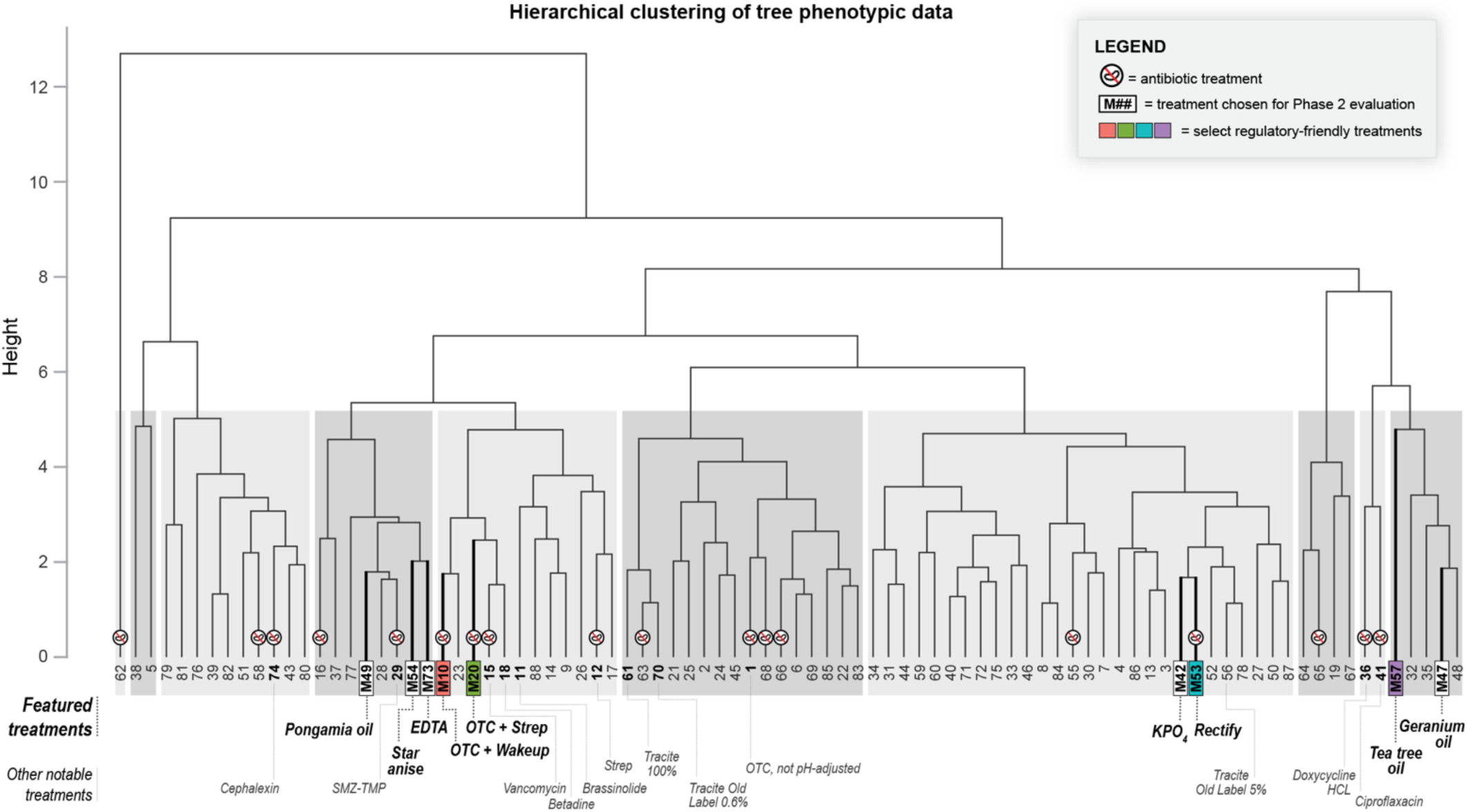
Hierarchical clustering of treatments based on multivariate phenotypic profiles. Mean values for ten traits were calculated for each treatment: average fruit weight, fruit number, soluble-solids content, change in tree health index from 0 to 180 days and 0 to 90 days, total fruit drop, pre-harvest canopy volume and canopy height, solids per box, and percent juice. Trait means were z-scaled, and pairwise Euclidean distances were computed, followed by agglomerative clustering with complete linkage. The dendrogram’s branch height reflects dissimilarity between treatments.

The fungicide propiconazole (M33), which was injected into the lowest quality trees (omega) also had a significant beneficial effect on the number of fruit >200 g produced (**Fig. 4a**, **Extended Data Table 7**).

### Retrospective power analysis of Phase 1 results

To inform experimental design for future grove-first screens, we performed a retrospective power analysis on the suite of data we collected in Phase 1. For reference, power curves for a one-sided t-test are presented in **Extended Data Fig. 11**. For this analysis, we selected a target false positive rate (alpha) of 0.05 and false negative rate (beta) of 0.05, representing a slightly lower tolerance for false negatives than a typical ‘standard’ criterion of alpha = 0.05, beta = 0.2. Under this ‘high power’ scenario, effect sizes of 3.3 times the standard deviation are required. **Extended Data Table 8** gives this effect size as a percent of the mean for the various parameters we measured in Phase 1, given the variance we observed in those parameters. Because variance was dependent on the class of tree observed, each combination of tree class and parameter is listed separately.

Detectable changes varied drastically by parameter and tree class observed, from as small as a 15% improvement in tree health index for alpha trees to as large as a 130% increase in fruit weight for omega trees (or a 94% increase for alpha trees). We also note that with all screens, there is an inherent tradeoff between sample replication and the number of treatments that can be tested. **Extended Data Table 8** also lists detectable changes for higher *n* and the requisite lower number treatments (e.g. *n* =6 and T=44 as opposed to *n* =3 and T=88). The outcome of common multiple hypothesis correction procedures on these data can be approximated by requiring a more stringent alpha in the power analysis (alpha = 0.001 in **Extended Data Fig. 11**). For small sample sizes, this scenario limits detection to prohibitively large effect sizes (∼9 times standard deviation for *n* = 3). This observation reinforces the absolute requirement for additional screening of candidate treatments from Phase 1, as our low replication, maximal treatment approach will generate false positive results that can only be eliminated by additional, Phase 2 experiments.

### Phase 2 analysis of selected chemistries

Midway through the 2023 growing season, a subset of Phase 1 treatments showing positive effects on foliar tree health were selected for evaluation in Phase 2 trials in commercial groves (**Figs. 2e, 3e**). A total of 13 treatments were evaluated Phase 2 analyses in three commercial citrus groves (**Extended Data Table 9**). These treatments included seven treatments from Phase 1, two treatment combinations from Phase 1, WakeUp alone and three proprietary iron-containing fertilizer formulations suggested by a grower (**Extended Data Table 9**). Regulatory status, (for example M10 would be immediately usable by growers, as surfactants added to tank mixes are not regulated by the EPA), as well as discussions with grower cooperators also heavily influenced our selection of treatments to evaluate in Phase 2.

In Phase 2, M10 received the highest priority for resources and trees because of the large positive foliar effects observed in the screen (**Fig. 1c, Extended Data Figs. 4, 6**) and that growers could use this treatment as a tank mix with no further regulatory actions. The trial site at Vero Beach West encompassed a row of 5-year-old, bearing Minneola trees treated with OTC (M53) and a row of M10 planted on the same bed (**Fig. 6a**). At 7-months post-injection, the canopy areas of the M10 treated trees were 39 times larger than the M53 treated trees (**Fig. 6b**, *P* = 0) and significantly increased the number of boxes per acre (Welch t-test *P* = 0.02, **Fig. 6c**). M10 treatment also significantly increased boxes per acre in Hamlin trees at the Hardee county site (Welch t-test *P* <0.0001, **Fig. 6c**). M10 had no significant effect on the pounds solids per box at any site when compared to OTC alone (**Fig. 6d**). A double injection of M10 spaced out six months apart in the Vero Beach West site resulted in the production of more and heavier fruit (**Extended Data Fig. 12**, ANOVA *P-*value <0.05), but internal quality was not significantly different as compared to the single M10 or M53 injections.

**Fig 6.**
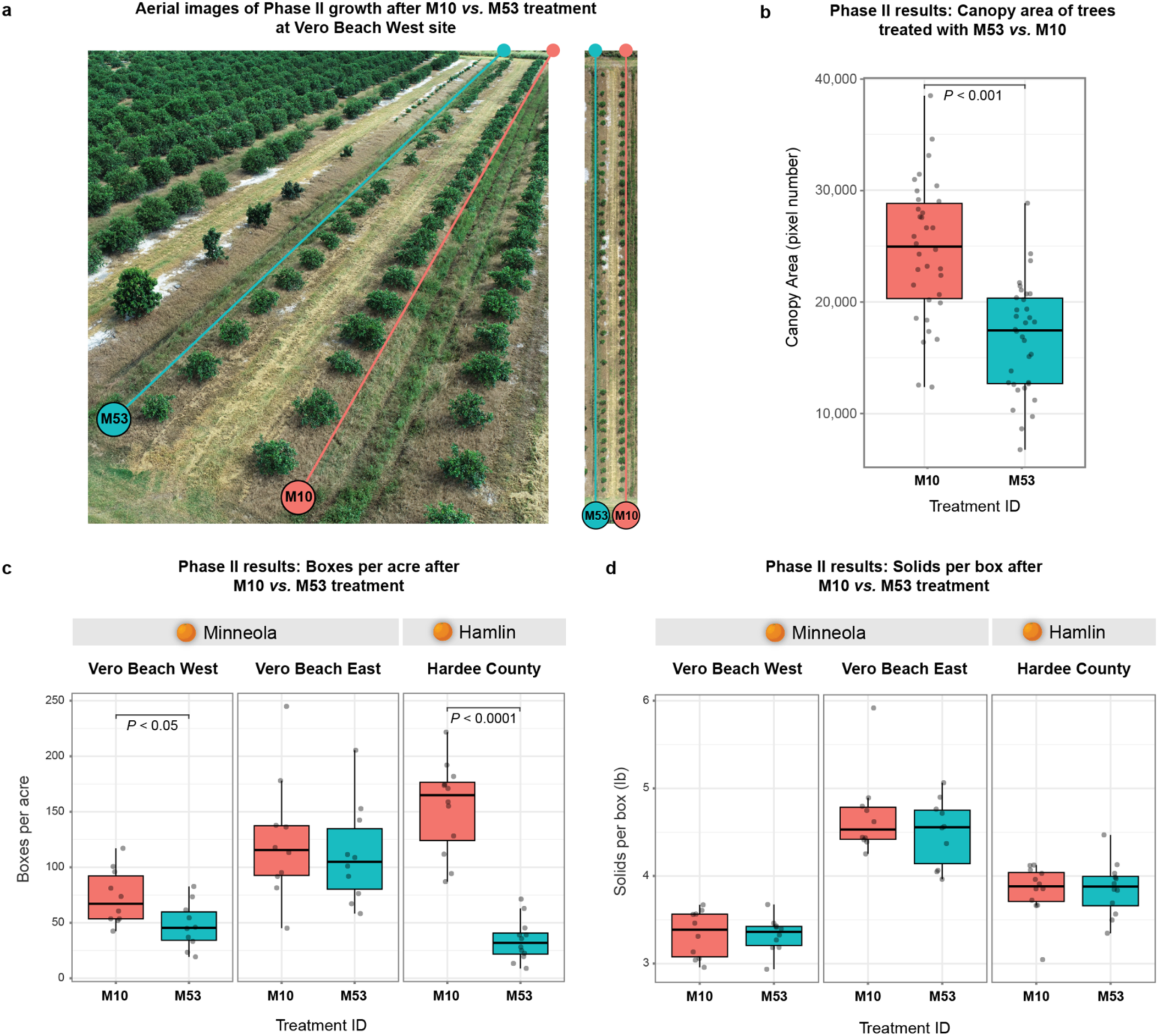
Phase 2 field performance of the oxytetracycline plus WakeUp treatment (M10) versus the oxytetracycline control (M53). (a) Drone photograph of the Vero Beach West grove seven months after injection, with the M10-treated row (red outline) showing visibly larger, denser canopies than the adjacent M53 control row (blue outline). (b) Distribution of individual-tree canopy areas at Vero Beach West (*n* = 30 per treatment), t-test *P-*value *=* 0; (c) Boxes-per-acre yield for Minneola and Hamlin blocks at three commercial sites; each panel depicts paired box-plots for M10 and M53 within the same orchard. Box per acre is significantly higher for M10 as compared to M53 at the Vero Beach West and Hardee County groves (Welch t-test *P-*value = 0.02 and *P*-value *=* <0.0001, respectively. (d) Pounds of soluble solids per field box recorded at the same sites and cultivar blocks, no statistically significant differences observed.

At the Vero Beach East test site, a total of eight treatments were evaluated. The grower requested the team to evaluate three of his own iron-containing formulations (referred to as FRC treatments) in addition to the Phase 1 selections. The grower also requested us to use the ReMedium TI^TM^ formulation of OTC for M53, which is a pharmaceutical grade product. The grower treatment FRC249 and combination of ReMedium TI^TM^ and WakeUp were statistically indistinguishable from ReMedium TI^TM^ alone across different tree metrics (**Extended Data Fig. 13**). Mixtures of geranium oil and WakeUp, clove oil and WakeUp as well as EDTA (M73), match ReMedium TI^TM^ for fruit weight, acidity and TSS to TA ratio, but have slightly lower °Brix (**Extended Data Fig. 13**). The grower’s other iron formulations FRC 11 and FRC 713 show either a mixed or inferior response as compared to ReMedium TI^TM^, respectively. The yield numbers at this site may be an under-representation of the true potential of these treatments because this grove sustained some damage during hurricane Milton in 2024, with the predominant hurricane-force winds hitting the east (injected) side of the trees.

At the Hardee County Phase 2 test site, a total of nine treatments were evaluated. OTC-WakeUp and potassium phosphate (M42) were clearly superior to M53, producing more boxes per acre, increased mean fruit weight, and maintained an equal or higher °Brix and soluble solids per box without increasing acidity (**Extended Data Fig. 14**). EDTA (M73), geranium oil (M47), pongamia oil (M49) and muriatic acid (M75) were statistically indistinguishable from Rectify^TM^ (M53) for all key traits (**Extended Data Fig. 14**). Clove oil (M59) raised acidity and doubled the produced boxes per acre, but delivered lower °Brix and the lowest TSS/TA ratio, therefore yields fruit of lesser flavor quality despite higher unit yield. WakeUp alone (no M#) showed no improvements as compared to Rectify^TM^ (M53) or other treatments.

## Discussion

The Grove-First framework departs from the standard lab to field pipeline and produced unanticipated positive results. Of the 88 chemistries initially tested, 13 were advanced to Phase 2 trials in a much shorter time frame than we anticipated with three new treatments producing horticultural responses on par with the currently available treatment, OTC-WakeUp (M10), potassium phosphate (M42) and FRC249. The focus of Grove-First to find any therapies that support the overall health of the trees under HLB pressure, not antimicrobial treatments specifically. While it has been demonstrated that OTC can both reduce *C*Las titers and improve fruit yield^30^, it has also been noted that there is generally poor correlation between *C*Las titer and fruit yield^31^.

Therefore, we devoted more time and resources to evaluating economically relevant outcomes in the largest number of trees possible, instead of measuring *C*Las abundance. Not making claims about antimicrobial activities of the different treatments that have a beneficial effect on tree health also allows them to be used by growers in manner consistent with EPA regulations, which relies on intent when determining if an agrichemical is FIFRA-exempt. Randomization of trees receiving treatments in the block was also important to our success, as the position in the block had effects on starting tree condition and yield metrics, independent of the treatment, including possible negative edge effects in row #8 and beneficial effects in row #1 closest to the *Murraya paniculata* hedge row. *M. paniculata*, a citrus relative, is known to be resistant to HLB and reduce *C*Las transmission by psyllids^32–34^.

The Grove-First approach integrates two key components to deliver effective therapies for managing HLB: drug repurposing for therapy selection and a design-of-experiments (DoE) framework for treatment testing in the field. Drug repurposing, a common approach in discoveries of treatments for human disease^35–37^, identifies new applications for existing therapies, leveraging billions of dollars of prior investment in regulatory, safety, supply chain, efficacy, and environmental testing. It dramatically reduces costs and expedites delivery compared to screening and developing new chemistries. Grove-First prioritizes therapies with established EPA regulatory profiles that are safe, affordable, and scalable, meaning the selection criteria we used considered regulatory alignment. For example, agricultural adjuvants, such as WakeUp, that may enhance product systemic movement and fertilizers, such as potassium phosphate or iron, are not regulated by the EPA. Some EPA categories are easier to repurpose, such as those exempting natural product oils and therapies that can be approved for citrus injection through label extensions, such as tea tree oil (M57). Tea tree oil is already registered by the EPA as a biochemical pesticide with fungicidal and bactericidal activity in a product called Timorex ACT (Summit Agro). Ongoing field trials with this labeled product will allow the company to determine if a label extension for citrus injection is warranted. DoE is a structured multivariate experimental approach widely used in industry to optimize and validate products and processes efficiently. Further, DoE ensures that clear and relevant objectives are defined, and all subject matter experts are actively involved throughout all stages of experimentation – planning, setup, data collection, analysis, and interpretation^4,5^.

Our results show a positive correlation between fruit weight and improved tree health metrics over the growing season, indicating that early selections for candidate treatments to advance into Phase 2 experiments can be made months prior to harvest, such as the WakeUp-OTC combination (M10). Similar early improvements in canopy density have been shown to correlate with increased fruit yield^31^. This early feedback is useful in the Grove-First screening framework because it allows lead time for regulatory adjustments to a product’s label if results support integration into HLB management. The focus on field outcomes over antimicrobial lab assays reinforces the idea that HLB is a disease complex^38^ and different treatments may be beneficial as part of a management plan that support the health of the tree and its entire microbial community.

An unexpected and positive outcome for the grower is the discovery that mixing OTC with other treatments, such as iron and surfactants, improved OTC efficacy. The Phase 1 to Phase 2 data collected for OTC and WakeUp (M10) in citrus varieties Valencia, Minneola and Hamlin demonstrated that the Grove-First DoE framework works as intended to identify treatments in the field with large effects. Combinations of iron and OTC also performed well in Phase I. These data highlight that there is room to improve the efficacy of an existing, EPA-labeled treatment, and work in this area is a high priority of research moving forward. The Phase 2 M10 results validated the framework we developed to identify treatments with large effects in a field screen by injection by showing the repeatability of a large phenotypic effect across multiple field sites.

A key component of Phase 2 trials is to validate a treatment effect in different environments to assess the robustness of the treatment. From the analysis of treatments evaluated in Phase 2, potassium phosphate (M42), a readily available fertilizer with no risk to the development of promoting antimicrobial resistance, is the most compelling non-antibiotic candidate to come from the study due to the repeatable benefits in Phase I and Phase 2. Future work in expanded Phase 2 or Phase 3 trials would provide further confirmation of the yield benefits.

The grower-identified FRC 249 showed an OTC-equivalent fruit quality in the Phase 2 trial. Potassium phosphate and FRC 249 did not rely on co-injections with any antibiotic, making them ideal as possible rotation partners if multi-site confirmation is obtained. FRC 249 is also a fertilizer and would not be regulated by EPA. Star anise and pongamia oils increased yield at Hardee County grove site, but would require flavor and consistency refinements before broader deployment. The Phase 2 results support a possible three-pronged approach for expanded field trials: integration of M10 (or other surfactants) for enhanced OTC action, nutrient supplementation with potassium phosphate (and iron) and a novel FRC candidate. Each of these treatments (M10, FRC, and potassium phosphate) were validated in one or more Phase 2 trials and hence would be ready for broader scale experimentation. Fertilizer and micronutrient support have previously been shown to improve the yield and tree health of HLB-infected citrus^39^. Collectively, these observations challenge the assumption that antimicrobial activity is the sole determinant of treatment efficacy and suggest that screening paradigms should incorporate broader physiological indicators - such as nutrient status and canopy density - to better predict field performance.

Phase 3 trials, beyond the scope of what we achieved here. These trials would include further ruggedness testing^5,40–42^, a DoE method to determine how robust a treatment is to varying operational conditions, such as variability in field crew techniques, timing of application, and environmental conditions after treatment. These are important factors to understand, as these can affect the treatment’s uptake, distribution, or degradation. In the U.S., the USDA Citrus Research and Field Trial Program (CRAFT) conducts ruggedness testing of EPA-approved therapies to examine how key factors influence the efficacy and reliability of a treatment, a treatment’s robustness, across different operational and environmental scenarios in all citrus-growing states.

We equally weighted regulatory status and tree performance metrics to advance treatments into Phase 2 evaluations and thus did not advance any new antibiotic treatments into Phase 2 trials in the first year, even though some antibiotics showed much larger effects on °Brix in the Phase 1 screen. This was a deliberate use of resources to focus on identifying injectable treatments that could be a non-antibiotic, rotational partner to OTC in the first year of screening. Ongoing Phase 2 trials in the field are incorporating other antibiotics in conjunction with residue testing to evaluate safety. The combination of OTC and streptomycin was previously shown to increase yield and fruit quality of Midsweet oranges by two °Brix points^18^. In that study, trees were injected four times every three months using 200mL injections, whereas we did a single injection using 100mL.

In our study, the °Brix increase for the OTC and streptomycin was less than 1 point, on average, but the increase in yield and production of large fruit was economically significant. Coupled to the Phase 2 results we had for the double M10 injections, these data collectively show that multiple injections per year is more efficacious than one. The sectoring we observed in the distribution of the foliar response may have resulted in an under-estimation of the °Brix values for some treatments, namely if the product was so unevenly distributed throughout the trunk of the tree so as to produce a visible skew in foliage response, it is reasonable to hypothesize that fruit from different sectors may have different ripening properties and quality profiles. These data suggest that growers may see enhanced effects from bi-lateral trunk injections. It is also possible that multi-year treatments are needed to see cumulative effects on improving tree health, as was demonstrated with two consecutive years of OTC injections^43^. Additionally, exploring strategies to make OTC more effective is also a priority, as the combination of WakeUp and OTC produced dramatic effects in the screen and Phase 2 evaluation. Given that the growers have armed themselves with a tool to deliver HLB therapies directly into the citrus vascular tissue, the Grove-First data show that the shortest path to overcoming the HLB crisis in Florida hinges on a commercial-scale demonstration that injectable treatments can solve the HLB problem economically. An injectable solution will enable the growers to buy time while the scientific community develops and deploys an engineered, HLB-resistant tree-of-the-future incorporating advances in biotechnology^44–48^.

## Materials and Methods

### Phase 1 Field Study

In 2023, the Phase 1 study was conducted at the United States Department of Agriculture, Agricultural Research Service (USDA-ARS) Picos farm located in Fort Pierce, Florida, USA (27°26′01.2″ N, 80°25′51.0″ W) using 8-year-old ‘Valencia’ sweet orange (*Citrus sinensis* L. Osbeck) grafted onto a US 812 (*Citrus reticulata* x *Poncirus trifoliata* L. Raf.) rootstock. They were 1.65 to 2.99m tall and 2.37 to 3.91m wide, planted at a density of 170 trees/acre. Trees received standard irrigation and fertilization care; however, no psyllid control was implemented. Prior to injections, tree trunk diameters 5 cm above the graft union were measured for each tree. The disease index rating method^49^ was modified to produce tree health ratings to classify trees into three different groups based on the trunk diameter and disease index rating: alpha, beta, or omega (**Extended Data Fig. 15, Extended Data Fig. 1**).

### Tree Injections

Trunk injections began on May 15, 2023, and ended on July 7, 2023. Injections were divided into nine sets over different dates, with a total of 88 treatments (**Extended Data Table 1**). Injections were made using the FlexInject^TM^ (TJ Biotech, LLC), following the manufacturer’s instructions. Holes were drilled to a depth of 25 mm approximately 5 cm above the graft union on the scion. The tip of the injector was then inserted directly into the drilled hole and the valve was opened to allow for tree uptake. Once all injectors were deployed in a set, uptake was monitored. After uptake, or seven days, injectors were removed from the trees. Details on injection volume, diluent and products can be found in **Extended Data Table 1**.

### Tree health monitoring

Visual ratings of tree health index and canopy density were used to assess changes in restoring tree horticultural traits. Ratings were taken before injections and at 90, 180, and 270 days after the injections. For tree health index ratings, the methodology described by Slinski, Page and Syvertsen (2016) was followed^49^. Each side of the tree was divided into four quadrants, and each quadrant was rated from 0 to 5, where 0 = no foliar disease symptoms visible, 1 = foliar disease symptoms on <20% of the quadrant, 2 = foliar disease symptoms on 20 to 40% of the quadrant, 3 = foliar disease symptoms on 40 to 60% of the quadrant, 4 = foliar disease symptoms on 60 to 80% of the quadrant, and 5 = foliar disease symptoms on >80% of the quadrant. The scores from each side of the tree were combined, resulting in a total possible tree health index ranging from 0 to 40. Canopy density was also rated on each side of the tree on a scale of 1 to 5, where 1 = very sparse, 2 = sparse, 3 = moderate, 4 = dense, and 5 = very dense^30^. The average rating from both sides determined the overall canopy density of the tree. In addition to tree health index and canopy density ratings, a set of seven pictures was taken per tree on the day of injection, and then at 30, 60, and 90 days after the injections. The photos consisted of three taken from each side of the tree and one at the injection site.

### Canopy Volume

The canopy volume was evaluated on the day of injections and on February 29, 2024. Each tree was measured for canopy height, canopy width parallel to the row, and canopy width perpendicular to the row. These measurements were then used to determine the canopy volume using the equation as reported in Obreza and Rouse (1993)^50^, where π = 3.14; TH = Canopy height and ACR = Average canopy radius.

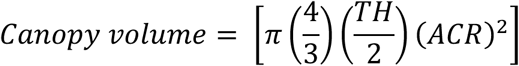

### Fruit measurements and harvest

Pre-harvest fruit drop was measured twice by counting and removing fruit on the ground beneath each tree on February 5, 2024, and March 8, 2024. The total number of fruits on each tree was counted at harvest and used to calculate the percentage of fruit drops for each tree. Harvesting took place on March 11, 2024. The total fruit weight per tree (kg per tree) was measured using a digital field scale (Ohaus Corporation, Parsippany, NJ, USA). Individual fruit were counted and weighed using a Compac 900CIR vision system (Compac, a Tomra Food company, Visalia, CA, USA).

Fruit juice content was determined using 20 fruits per tree, weighed, and extracted using a mechanical juice extractor (model 2702; Brown International Corp., Covina, CA, USA). The juice content was weighed to determine the juice percentage from Gottwald et al. 2012^22^:

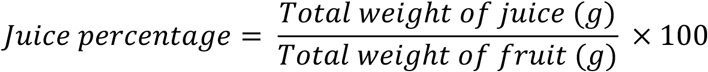

Soluble solids were measured using a refractometer (HI96801; Hanna Instruments, Woonsocket, RI). A pipette was used to deliver two to three drops of juice in the clean glass measuring surface of the refractometer. Fruit acidity was measured with an automatic potentiometric titrator (HI931; Hanna Instruments, Woonsocket, RI). For titration, 25 mL of sample juice was diluted with deionized water to make 50 mL. The solution was later titrated using a 1N sodium hydroxide solution. Fruit quality was also expressed as the ratio of soluble solids content and titratable acidity. The amount of pound solids is the soluble solids (sugars and acid) contained in one box of citrus fruit and was calculated using the formulas from (Wardowski et al. 1995)^51^:

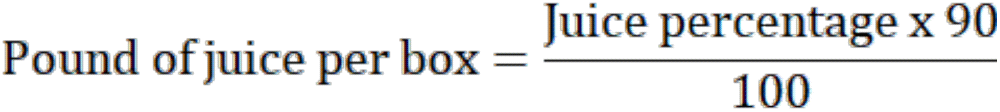

Where 90 is the weight of a box of citrus fruits in pounds.

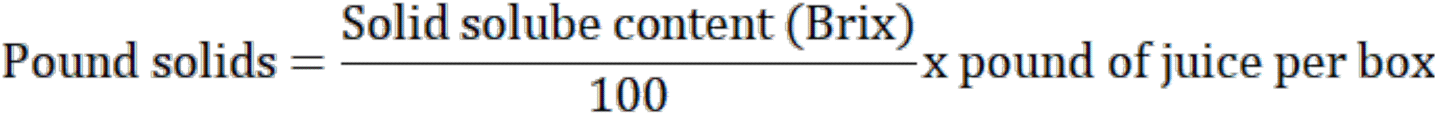

### Oxytetracycline residue testing

On January 25, 2024, twelve fruits, three from each of the cardinal directions (N, S, E, & W), were randomly selected from each tree in treatments containing oxytetracycline (M1, M10, M20, M58, & M68). These fruits were weighed and sent to the USDA AMS S&T LATD National Science Laboratories in North Carolina for oxytetracycline residue testing.

### Bacterial growth inhibition plate assays

A total of 87 out of the 88 treatments (we did not test the water control) were tested for antimicrobial activity against *Agrobacterium tumefaciens* strain EHA105. Initially, a pure culture of a single colony of EHA105 was inoculated into 5 mL Luria Broth (LB) medium in a tube and incubated in an orbital shaker (200 rpm) at 28°C overnight for 24 hr. Subsequently, a subculture was prepared from the overnight culture by diluting it with fresh LB medium to an optical density (OD) of 0.5 at 600 nm. 100 μL aliquots were added into tubes that contained 100 μL of test compounds at a 1% concentration. The mix was incubated in an orbital shaker (200 rpm) at 28°C for 1 hr and then plated onto Agar medium at 28°C for 24 hr. Bacterial growth was visually assessed and recorded for growth or growth inhibition (**Extended Data Table 1**).

### Plant material and tree injections for Phase 2 trials

The Phase 2 injections were carried out in collaboration with three citrus growers in Florida. Some of the treatments tested in Phase 1 that showed good performance were advanced to Phase 2, along with treatments submitted by growers, to be tested for what was called the “down-the-row effect.” The principle of the “down-the-row effect” was that the treatments were injected into a full row or half a row to make the effects of the injections more visible, using the Flexinjetor^TM^ (TJ Biotech LLC), following the manufacturer’s instructions. In all the fields tested the treatment M53 was used as a control. At the Hardee County site, *Citrus sinensis* (Hamlin) were used, but the grower did not know the rootstock. Between 22-28 trees were injected for each treatment and a total of 13 randomly selected trees (age-matched) for each treatment were sampled for yield and quality analysis, as described above. At the Vero Beach East site, mature Minneola trees were used on Kinkoji rootstock. A total of 20 trees were injected and a random selection of 10 trees per treatment were sampled for yield and quality analysis, as described above. At the Vero Beach West site, a total of 35 Minneola trees on UFR17 rootstock (planted 12/4/19) were injected with M53 and M10 in the bed designated in Fig. 6. A total of 168 trees were injected with M10 at two time points in the growing season. A total of ten trees were sampled for yield and juice quality, as described above.

### Drone imaging of Vero Beach West grove site Phase 2 trial

Aerial images were captured with a DJI Phantom 4 Pro drone (1-inch 20MP CMOS sensor) in sRGB color, 3:2 format, at 5472 x 3648 pixels. Photos were taken automatically at 46 meters altitude and 12 kph, using a straight-down orientation (-90° camera angle) to the trees, with 70% side-overlap and 80% forward-overlap to reduce relief displacement. Images were analyzed in ImageJ^52^, where tree canopy area was measured by counting the segmented pixels representing each tree.

### Data analysis and visualization

RStudio was utilized for statistical analyses and visualization^53^. All data and code described in this paper for the Phase 1 screen can be found on GitHub (https://github.com/MichelleHeck77/Grove-First). To compute the change in plant health index over time, plant-health scores (*n* = 264 trees) were imported from an Excel worksheet into R. Scores of “10” (indicating a moribund canopy) were re-coded to “5” to place all ratings on a 0 to 5 scale. For each tree, we calculated the baseline (0 day) health of four quadrants on the east and west sides of the canopy and then subtracted the corresponding values at 90, 180, and 270 days to obtain quadrant-level tree health change scores. These quadrant values were summed to obtain side-level totals (East0, West0, etc.), which were then combined to give whole-tree totals (Tree0, Tree90, Tree180, Tree270, Tree450). Finally, tree-level changes in health (Tree_delta_90, Tree_delta_180, Tree_delta_270 and Tree_delta_450) were computed as the difference between the baseline total and each subsequent time-point; these delta-values were used in all downstream analyses. For treatment-level pattern discovery using a correlation analysis (**Fig. 5**), we aggregated the full dataset by *MoleculeID* in R. Mean values of 10 quantitative phenotypes, including average fruit weight, fruit number, °Brix, tree-health change scores (90 day and 180 day), total fruit drop, pre-harvest canopy volume and height, box solids, and percent juice, were computed for each treatment. The resulting matrix (one row per molecule) was auto-scaled to account for the diversity of measurement units, and pairwise Pearson correlations were inspected to verify data structure. Hierarchical clustering was performed on the scaled Euclidean distance matrix using complete-linkage agglomeration^54^. The dendrogram was visualized and arbitrarily cut at *k = 10* to define ten discrete molecule clusters for downstream comparative analyses. There are any number of ways to perform hierarchical clustering and the code is provided to facilitate users’ ability to test different permutations (https://github.com/MichelleHeck77/Grove-First). These phenotypes were selected as they were the ones most meaningful to the growers. We made bubble plots that summarize treatment-level responses (Figs. 2 and 3). For each candidate chemistry in a particular category, we averaged six key measurements: tree-health change at 180 days, fruit-drop count, pounds of solids per box, °Brix, juice volume, and the TSS:TA ratio. Two plot types were generated: one shows tree-health change (x-axis) against fruit drop (y-axis) with bubble size indicating solids per box, and the other shows °Brix (x-axis) against the TSS:TA ratio (y-axis) with bubble size reflecting juice volume. Power curves (**Extended Data Fig. 11**) were generated in python with the statsmodels package, version 0.13.5. Phase 1 data were largely analyzed using Welch’s ANOVA and/or followed by t-tests to account for non-homogeneous variance and for comparisons to M53 or to tree category means (**Extended Data Tables 2-7**). Phase 2 data were largely analyzed using ANOVA at confidence level 0.95, alpha = 0.05, and the Tukey method was used for *P*-value adjustment. Data were checked for normality and homogeneous variance.

## Extended Data Figures and Legends

**Extended Data Fig. 1.**
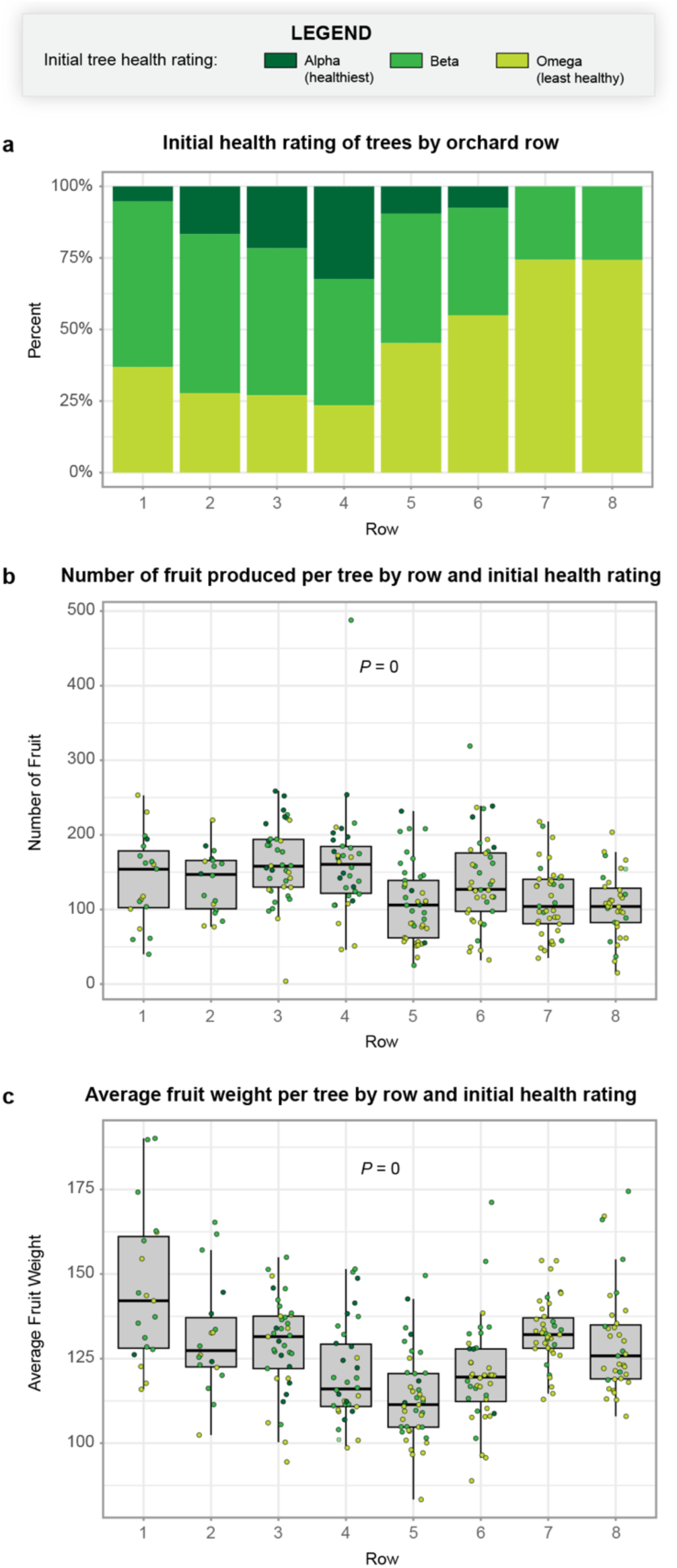
Influence of grove row and tree-health rating on yield components. (a) Proportional distribution of trees in each health rating class for every grove row. Bar height represents the percentage of trees in a given row that fall into the indicated rating category. The distribution of tree-health ratings differed across rows (Pearson’s χ² = 55.5, Monte-Carlo *P* = 0.0001, *n* = 264). (b) Total number of fruit harvested from individual trees as a function of grove row. Jittered points (one per tree) are colored by tree rating, matching the legend in (a). (c) Average fruit weight (g) per tree across grove rows, displayed as in (b). (b) The number of fruit per tree varied among rows (Kruskal-Wallis *P* = 0), and (c) average fruit weight showed an even stronger row effect (Kruskal-Wallis *P =* 0). Each point represents one tree.

**Extended Data Fig. 2.**
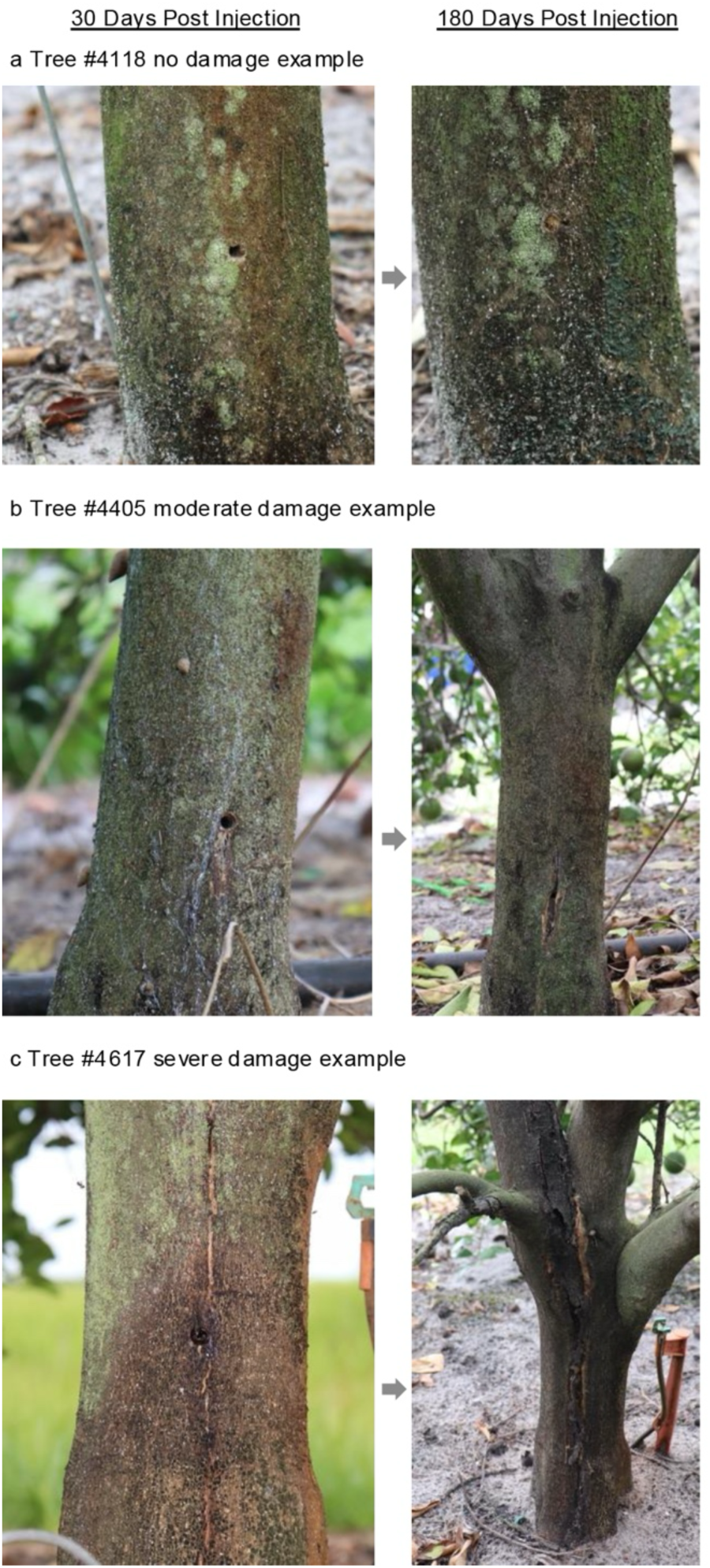
Representative spectrum of citrus trunk responses to therapeutic injection over six months. Paired photographs show the same injection site 30 days post-injection (left column) and 180 days post-injection (right column). (a) Tree #4118 (no visible damage) displays only the original injection site at 30 days and a fully occluded wound with no external lesions at 180 days. (b) Tree #4405 (moderate damage) shows a narrow vertical fissure and mild bark discoloration that persist and extend slightly after six months. (c) Tree #4617 (severe damage) develops extensive longitudinal cracking, necrotic tissue, and dark exudate radiating from the injection site, injuries which intensify over time.

**Extended Data Fig. 3.**
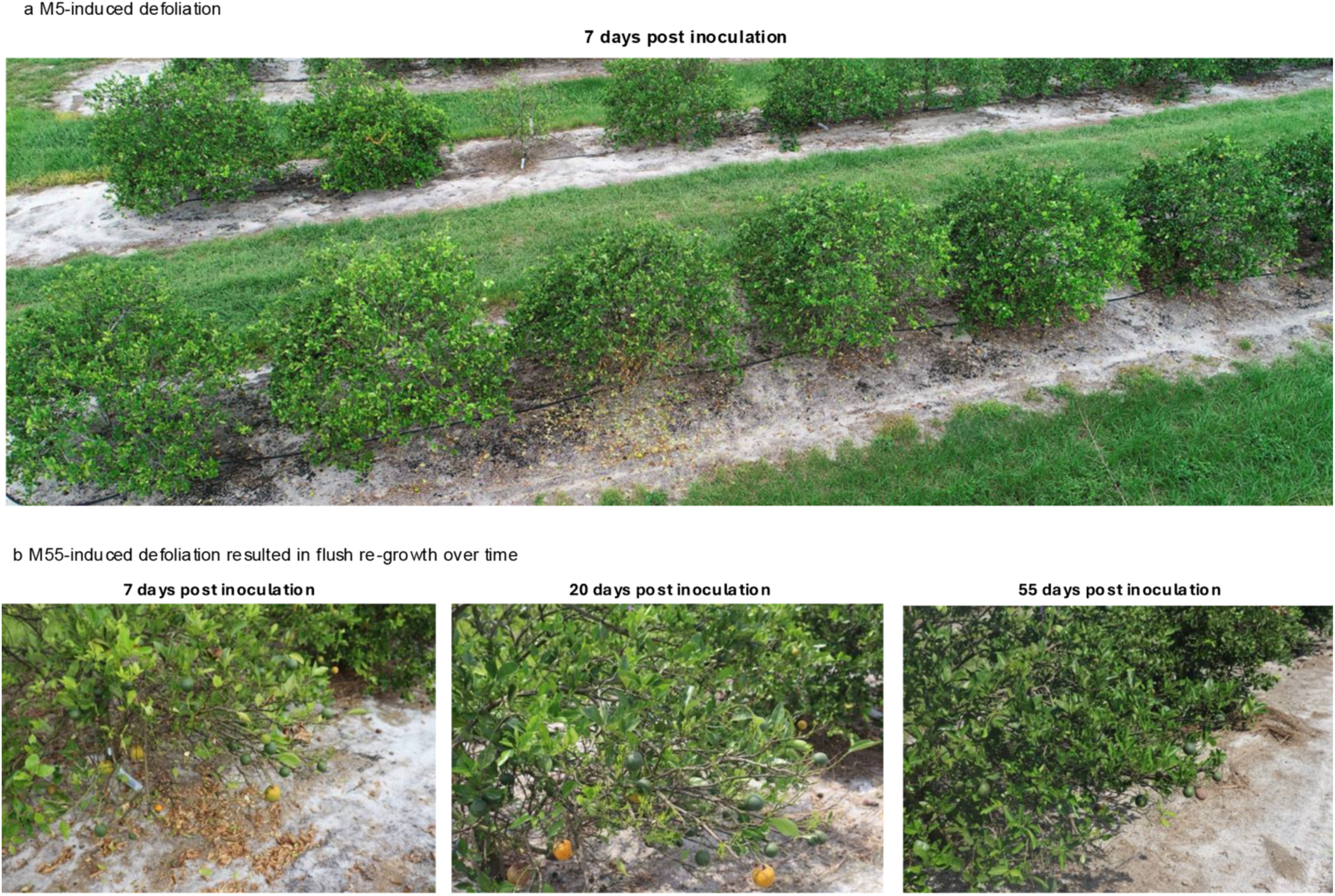
Localized, transient defoliation and canopy rejuvenation observed for some treatments following trunk-injection. (a) Grove overview 7 days post-injection (dpi) illustrating that the Instill^TM^ fungicide and bactericide (M5) treatment induced leaf abscission only on the flank where the injector was inserted (foreground side of the row); the opposite, non-treated side retained a normal canopy. (b) Time-course photographs of a representative tree taken at 7, 20, and 55 dpi (left-to-right) injected with Rectify^TM^ OTC plus Tracite®, an iron and sulfur chelated micronutrient solution (M55). The initial treatment-induced defoliation recorded at 7 dpi is followed by vigorous flush growth beginning by 20 dpi and culminating in a noticeably fuller, healthier canopy on the injected side by 55 dpi, surpassing the density of foliage on untreated portions of the tree. The sequence demonstrates that M55-triggered defoliation is reversible and can stimulate robust regenerative growth rather than long-term damage.

**Extended Data Fig. 4.**
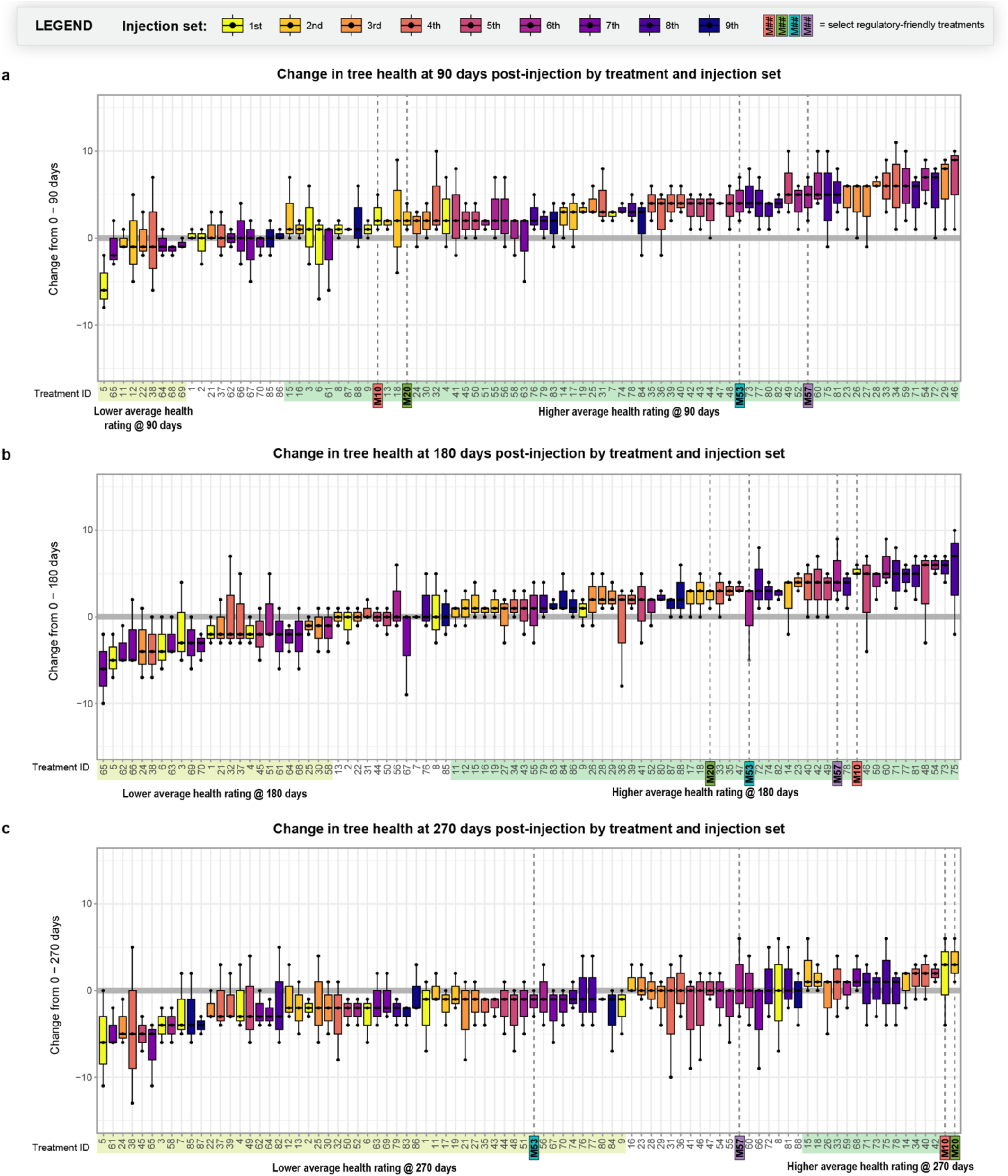
Effect of trunk-injected test molecules on citrus plant health index over three survey intervals. Box plots summarize the change in whole-tree plant health index (baseline Day 0 minus score at the indicated day = change in plant health index) for every experimental tree assigned to each treatment. Treatments (x-axis) are ordered within each panel by the median response. Each dot represents the score for each of the three trees in the screen per treatment (*n* = 264). Colors identify the sequential injection sets in which the treatments were deployed (legend above the panels); the teal symbol marks the suite of selected treatments discussed in Figs 4 and 5. (a) Acute response 0 to 90 days post-injection (dpi); (b) Mid-term response 0 to 180 dpi; and (c) Extended response 0 to 270 dpi. For all panels, the horizontal red line at change in plant health index = 0 denotes no net change in canopy condition over time. Values above the line indicate tree health improvement, whereas negative values indicate decline. Injection set 1-9 is denoted by color to show that there was no bias resulting from injection set.

**Extended Data Fig. 5.**
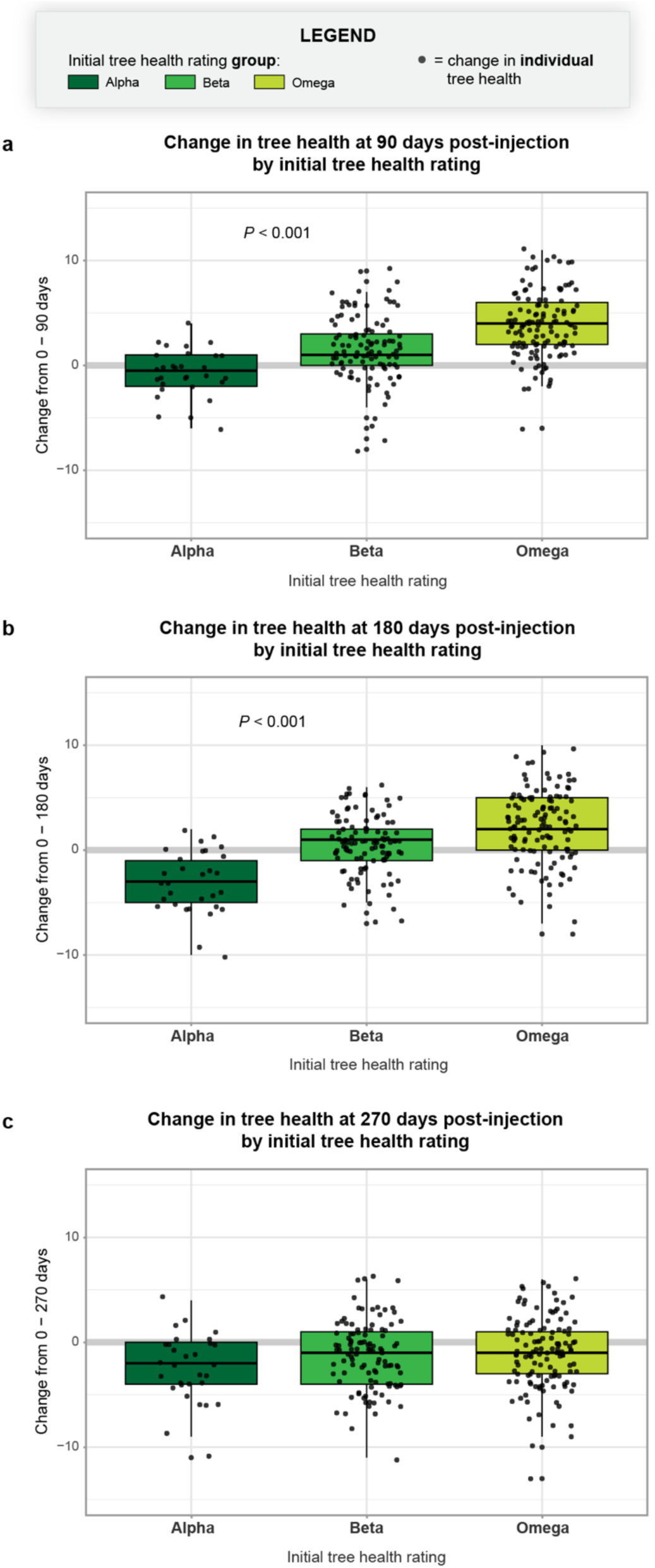
Baseline canopy condition predicts the magnitude and persistence of treatment-induced changes in plant health index. Boxplots show the distribution of whole-tree plant health index change for trees grouped by their pretreatment visual health rating (Tree Rating: “alpha” = healthiest, progressing to “omega” = most HLB-affected). Points represent individual trees. (a) Early response 0 to 90 days post-injection (dpi); (b) Mid-term response 0 to 180 dpi; and (c) Extended response 0 to 270 dpi. Across the two earlier time intervals, omega trees in poorer initial condition exhibited larger positive changes in plant health index, i.e., greater symptomatic recovery (Welch’s ANOVA *P-*values < 0.001 for 90 and 180 dpi, *P*-value at 270 days was n.s.).

**Extended Data Fig. 6.**
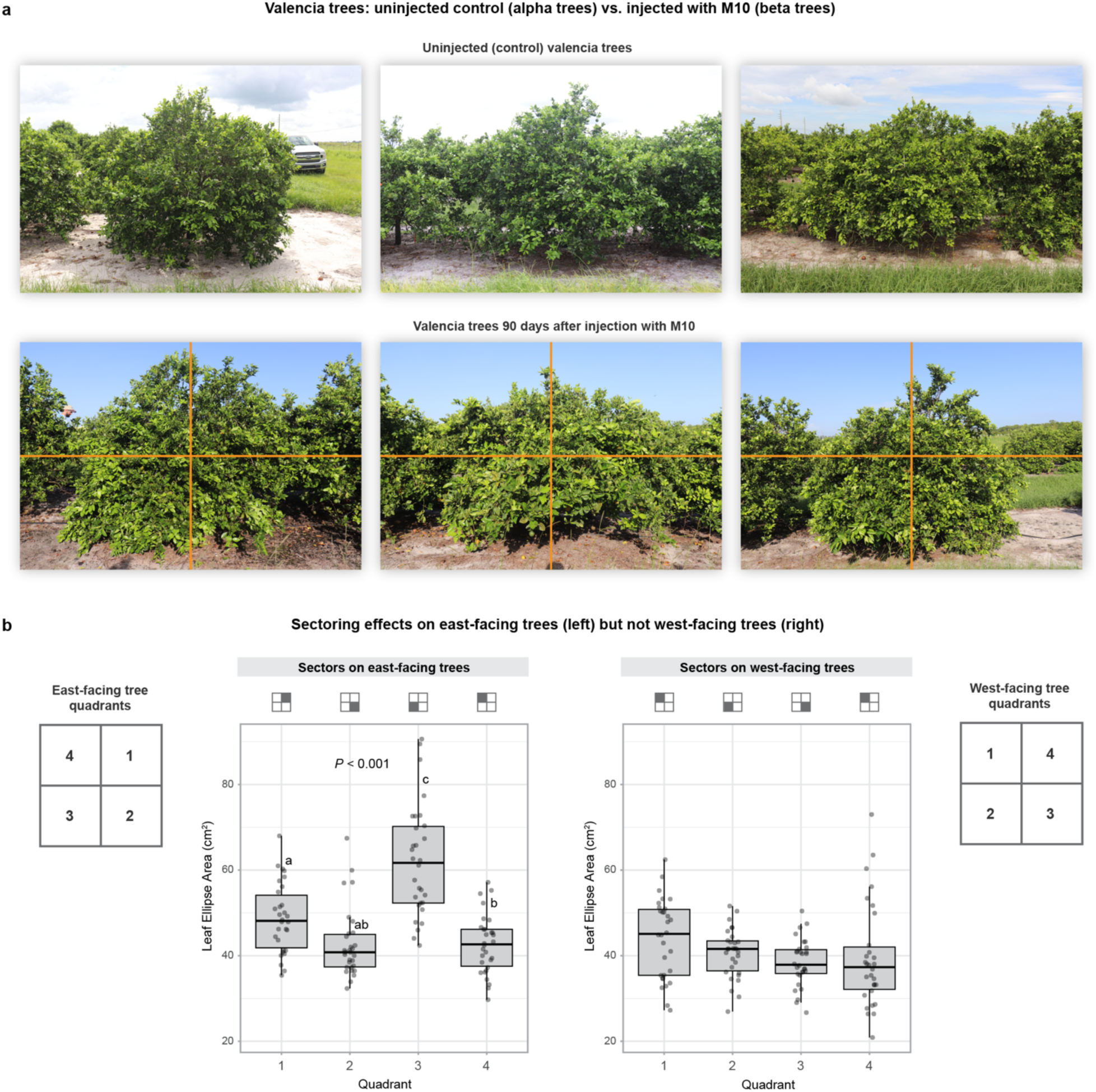
Sector-specific canopy responses reveal limited within-tree movement of some trunk-injected compounds. (a) Untreated trees (upper row) show a uniformly small-leaf canopy, whereas representative trees that received M10 (second row) develop conspicuously enlarged leaves only in discrete portions (“sectors”) of the crown. Vertical and horizontal orange guide-lines delineate the quadrant grid (Q1 - Q4) that was used for quantitative scoring of every tree side (inset diagrams at far left and far right). (b) Leaf size (leaf-ellipse area) depends on both the canopy side and quadrant. A two-way ANOVA detected main effects of east (injected side) vs west (*P* <0.001), quadrant (*P* <0.001) and a strong east/west × quadrant interaction (*P<*0.001). Tukey-adjusted pairwise comparisons showed that on the east side the upper-inner quadrant (Q3) had the largest leaves, Q1 was intermediate, and Q2 and Q4 were smallest; on the west side, leaf size did not differ among quadrants.

**Extended Data Fig. 7.**
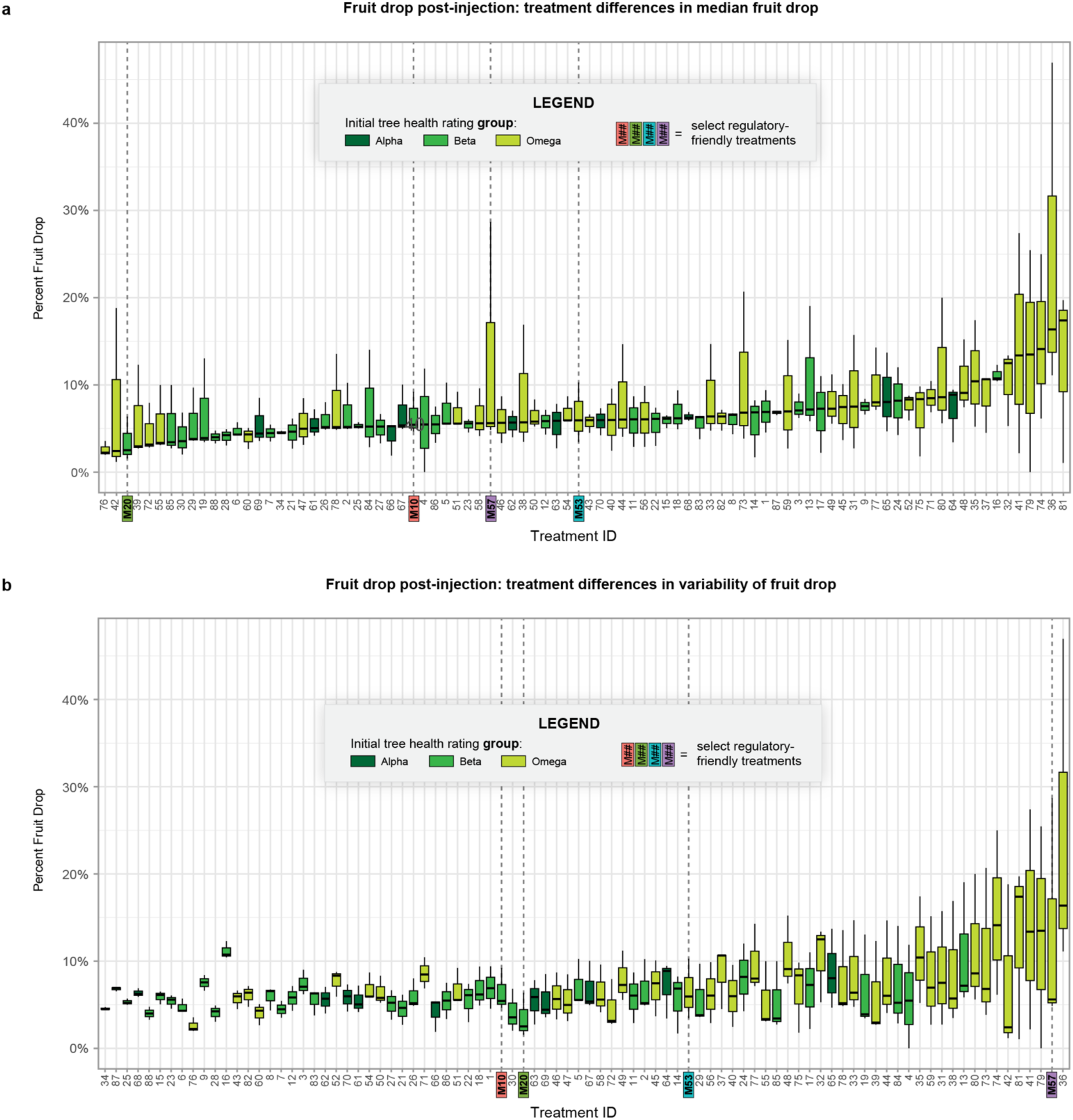
Premature fruit drop varies among test treatments and interacts with baseline tree health status. Box plots depict the percentage of fruit that dropped from individual trees prior to harvest for every experimental treatment (x-axis). Fill color denotes the initial tree rating class (“alpha” = healthiest through “omega” = most HLB-affected), enabling visual assessment of initial tree rating-dependent effects. (a) Treatments ordered by their median fruit-drop percentage highlight compounds that consistently minimize fruit drop (left side) versus those associated with elevated drop (right side). (b) Treatments reordered by the within-treatment standard deviation of fruit drop illustrate variability in response; wider spreads identify compounds whose performance is highly tree-specific, whereas narrow spreads indicate uniform efficacy (or lack thereof) across health classes.

**Extended Data Fig. 8.**
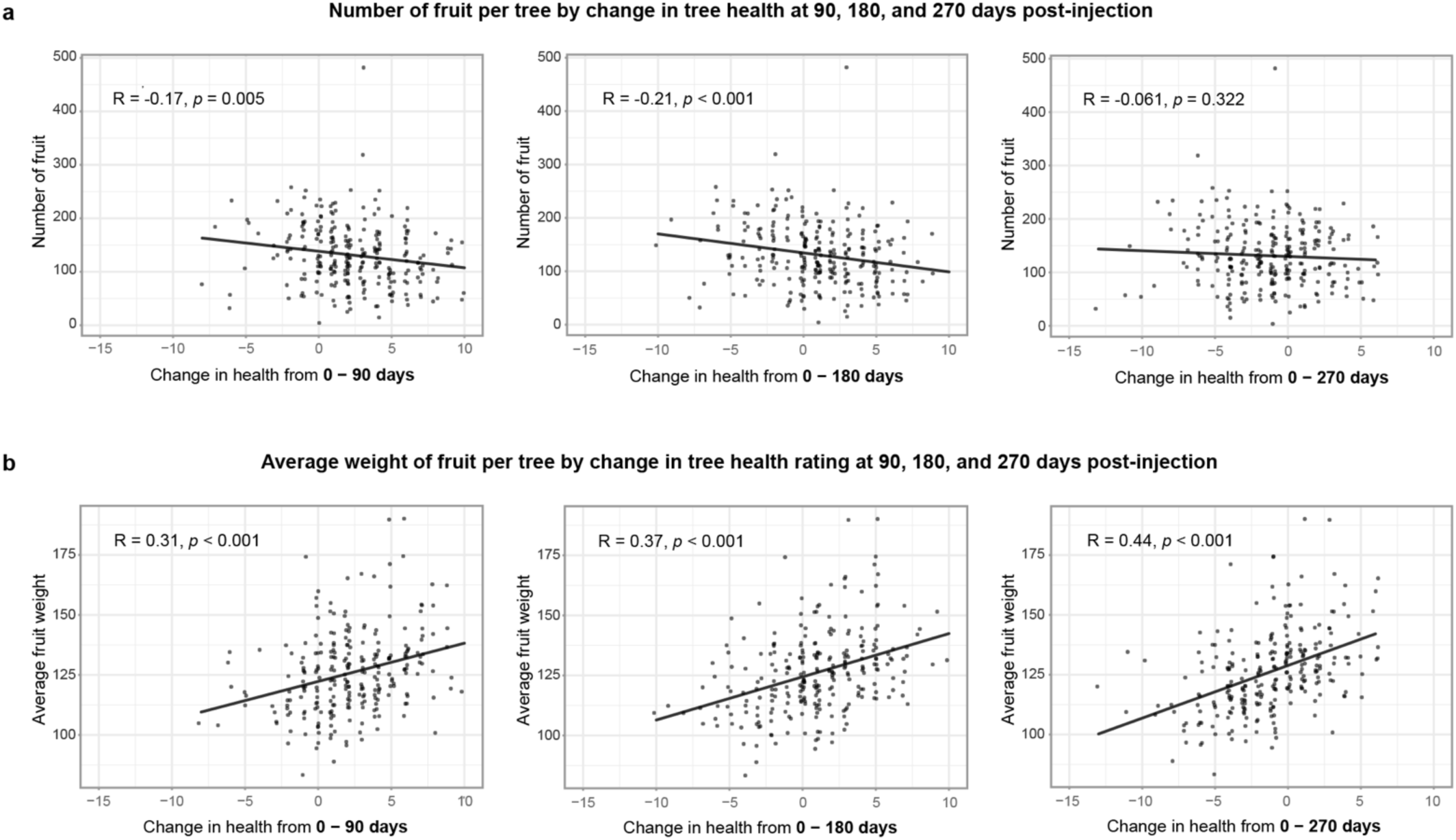
Early changes in canopy health differentially influence fruit load and individual fruit mass. (a) Number of fruit per tree. At 90 and 180 days post injection (dpi) there is a slight negative correlation: trees showing the largest visual health gains tended to carry fewer fruit, likely reflecting an initial trade-off between recovery and reproductive output. By 270 dpi, the relationship is not significant. (b) Average weight of each fruit per tree (in grams). In contrast, the change in plant health index is positively correlated with fruit weight at every interval. Trees that improved most produced larger individual fruit throughout the season. The Pearson correlation coefficients (R) and associated *P*-values are given in the panels.

**Extended Data Fig. 9.**
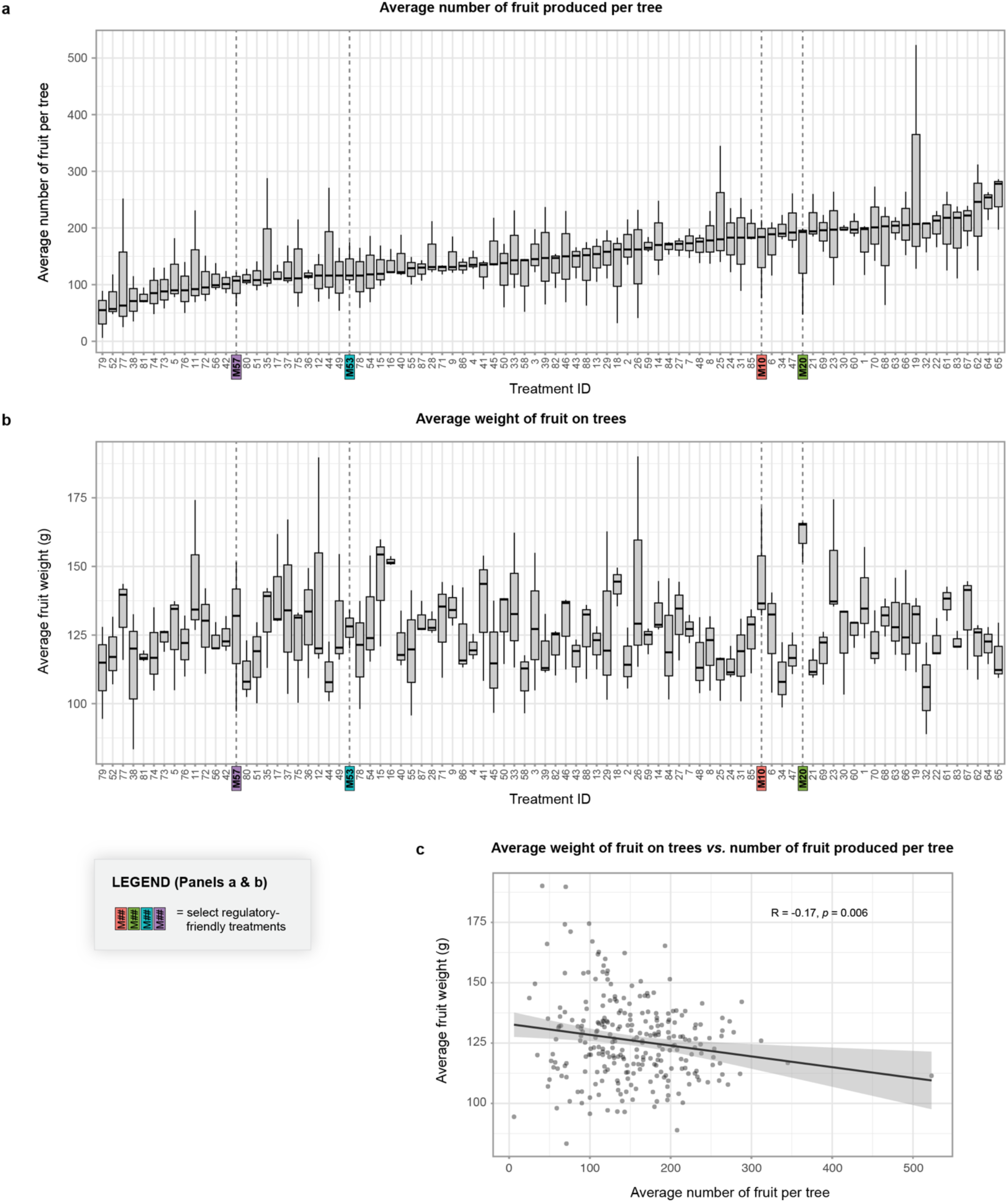
Treatment-level effects on fruit load, average fruit weight, and the trade-off between the two yield components. Total fruit, which is the sum of fruit captured by the Compac sorter, fruit dropped to the ground, and fruit removed from skirted branches prior to harvest is used for this analysis. (a) Total fruit per tree. Plots show the distribution of total fruit counts for every injected treatment, ordered by the median value. (b) Average fruit weight per tree. Box plots display the mean mass of individual fruit harvested from the same trees, using the identical treatment order as in (a) for direct comparison of load and size responses. (c) Fruit load versus fruit size across all trees. Scatter plot of total fruit count (x-axis) against average fruit weight (y-axis) with a least-squares regression line (black). The Pearson correlation coefficient (R) and associated *P*-value are given in the panel. The negative slope confirms a load-size trade-off, independent of treatment.

**Extended Data Fig. 10.**
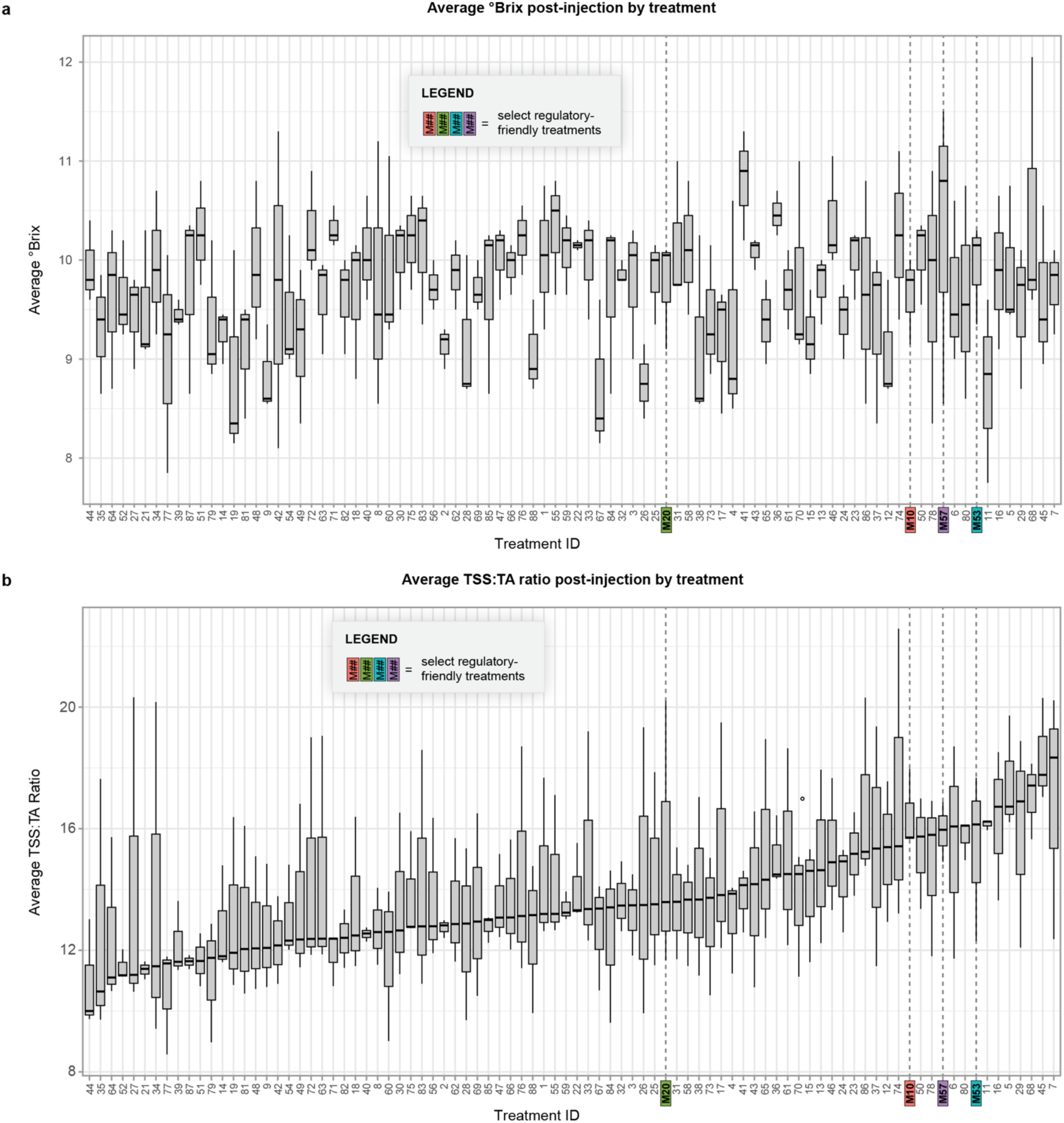
Soluble-solids accumulation and the sweetness-to-acidity balance in fruit harvested from each experimental tree across the 88 treatments in Phase 1. Treatments (x-axis) are ordered left-to-right by their median TSS:TA ratio so that panels (a) and (b) share an identical treatment order for direct visual comparison. (a) Average °Brix. Mean total soluble-solids concentration (°Brix) of expressed juice from each tree. Higher values denote greater sugar content. (b) TSS:TA ratio. Ratio of total soluble solids (TSS) to titratable acidity (TA), a standard industry metric of perceived sweetness relative to sourness. Larger ratios correspond to a sweeter flavor profile.

**Extended Data Fig. 11.**
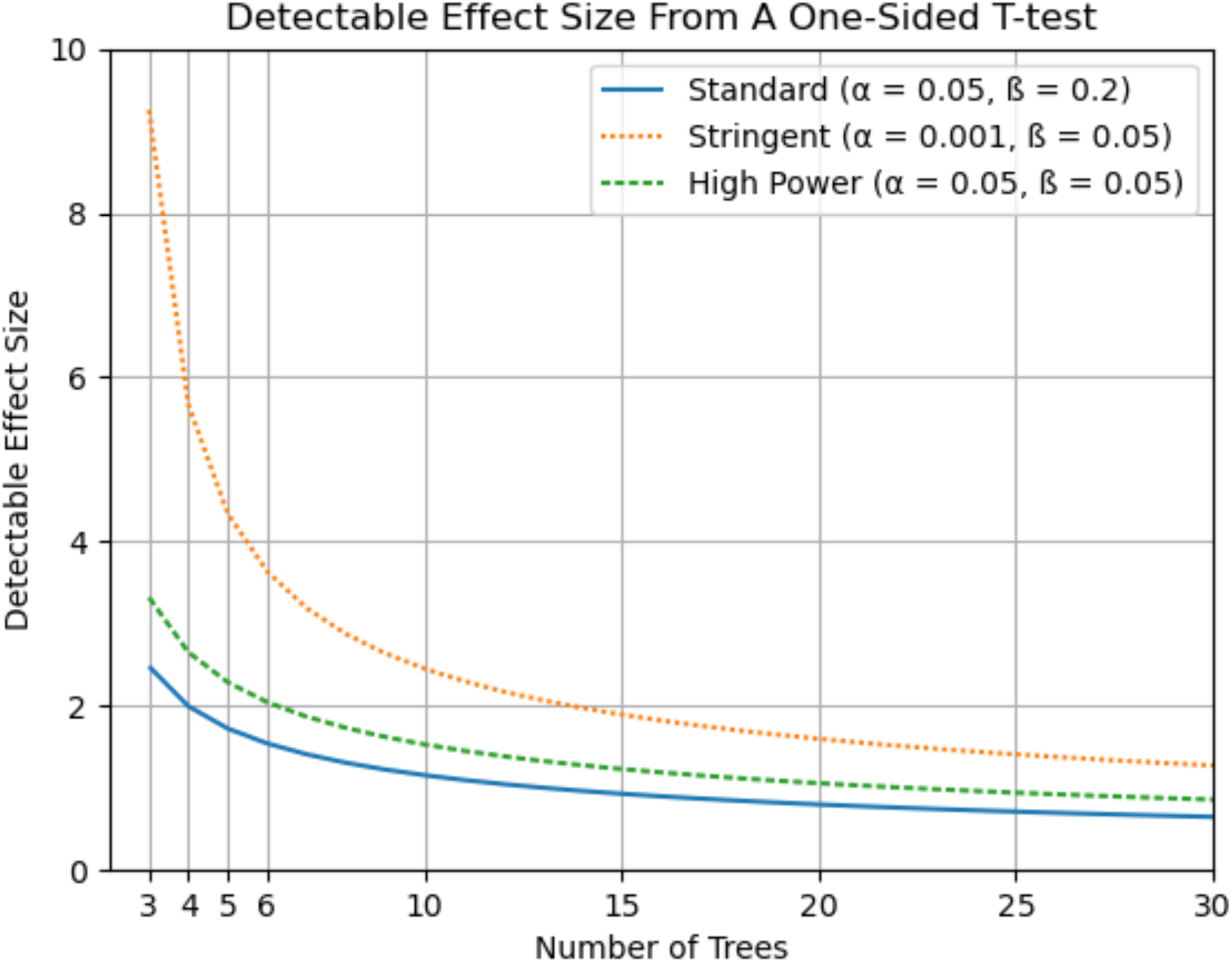
Power curves for a one-sided t-test under different false positive (alpha) or false negative (beta) rates. Detectable effect size (difference in mean divided by standard deviation) plotted against number of injected trees per treatment.

**Extended Data Fig. 12.**
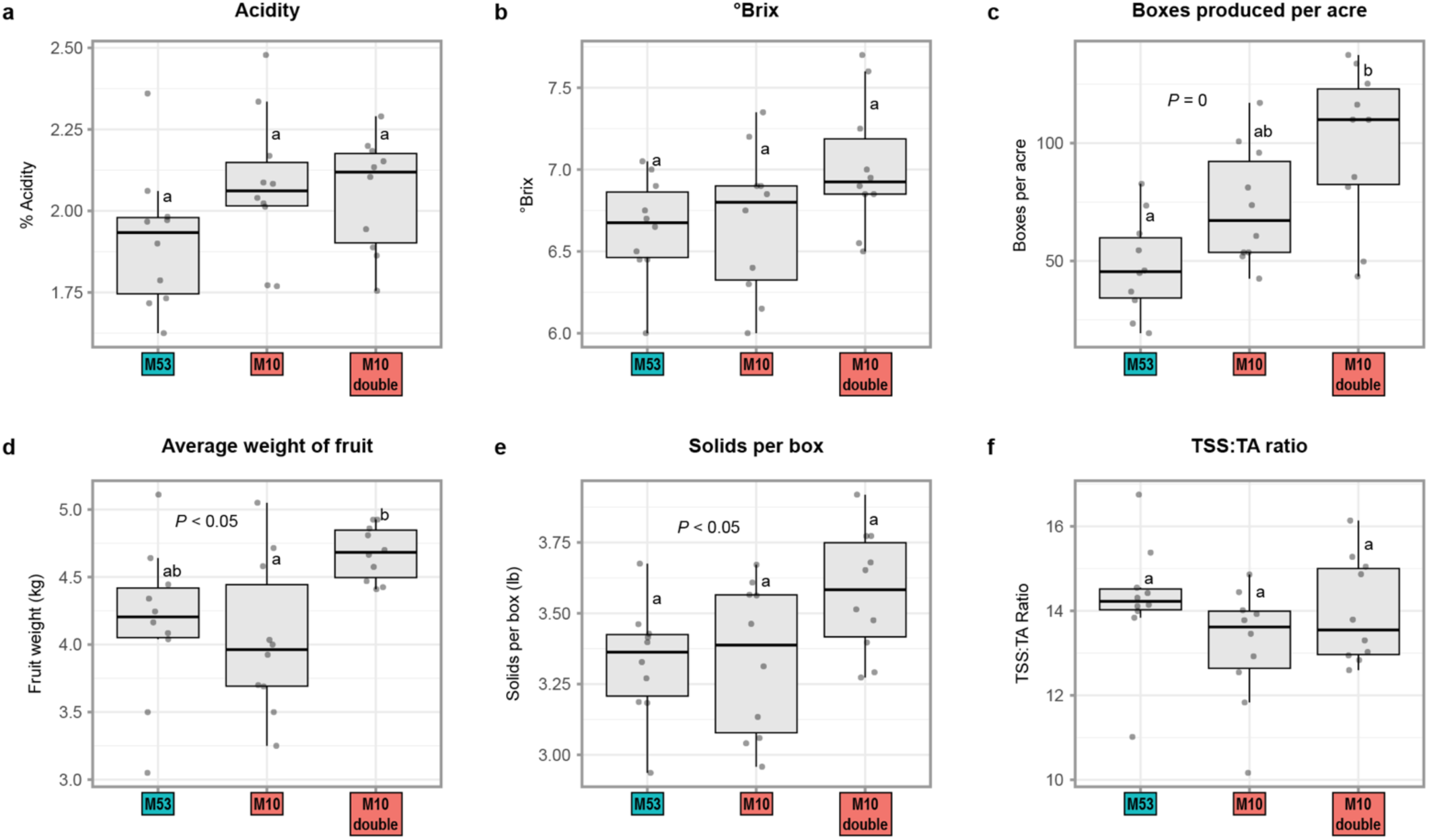
Phase 2 field performance of M10 in the Vero Beach West grove site. Traits include total boxes per acre, mean fruit weight, °Brix, titratable acidity, solids per box, and the TSS:TA sweetness-to-acidity ratio. Both M10 regimens (single or double injection) elevate yield components; the double dose delivers the largest gains in box yield and fruit weight but slightly lowers °Brix compared with the single dose, resulting in a comparable overall TSS:TA ratio. 5-year-old Minneola trees (planted 12/4/19) were used on UFR17 rootstock.

**Extended Data Fig. 13.**
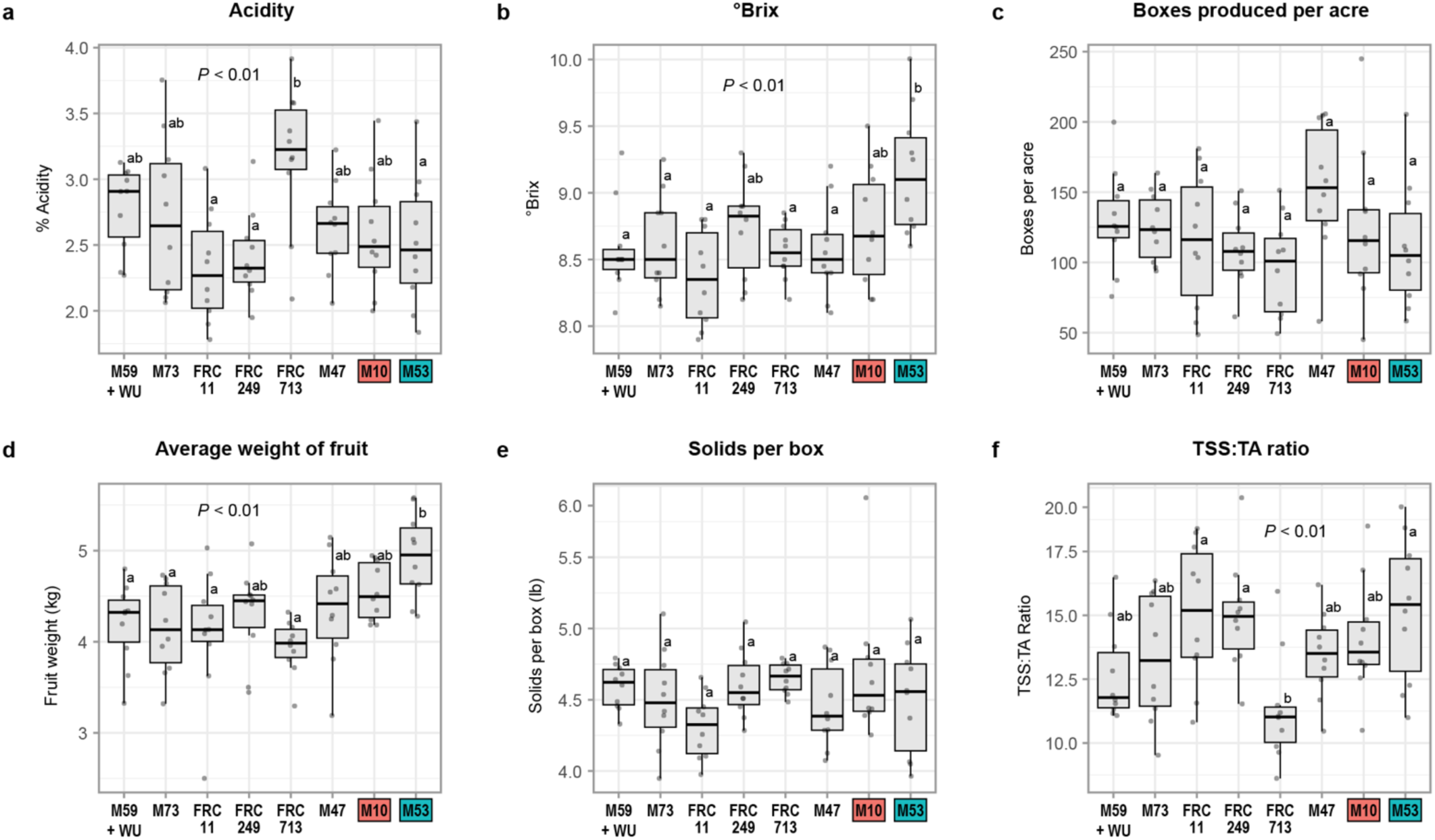
Phase 2 field performance of selected Phase 1 chemistries in Vero Beach East grove site. Traits include total boxes per acre, mean fruit weight, °Brix, titratable acidity, solids per box, and the TSS:TA sweetness-to-acidity ratio for selected treatments as indicated. Grower supplied treatments labeled as FRC#, which were formulations of a proprietary iron micronutrient compound. Mature Minneola trees were used on Kinkoji rootstock.

**Extended Data Fig. 14.**
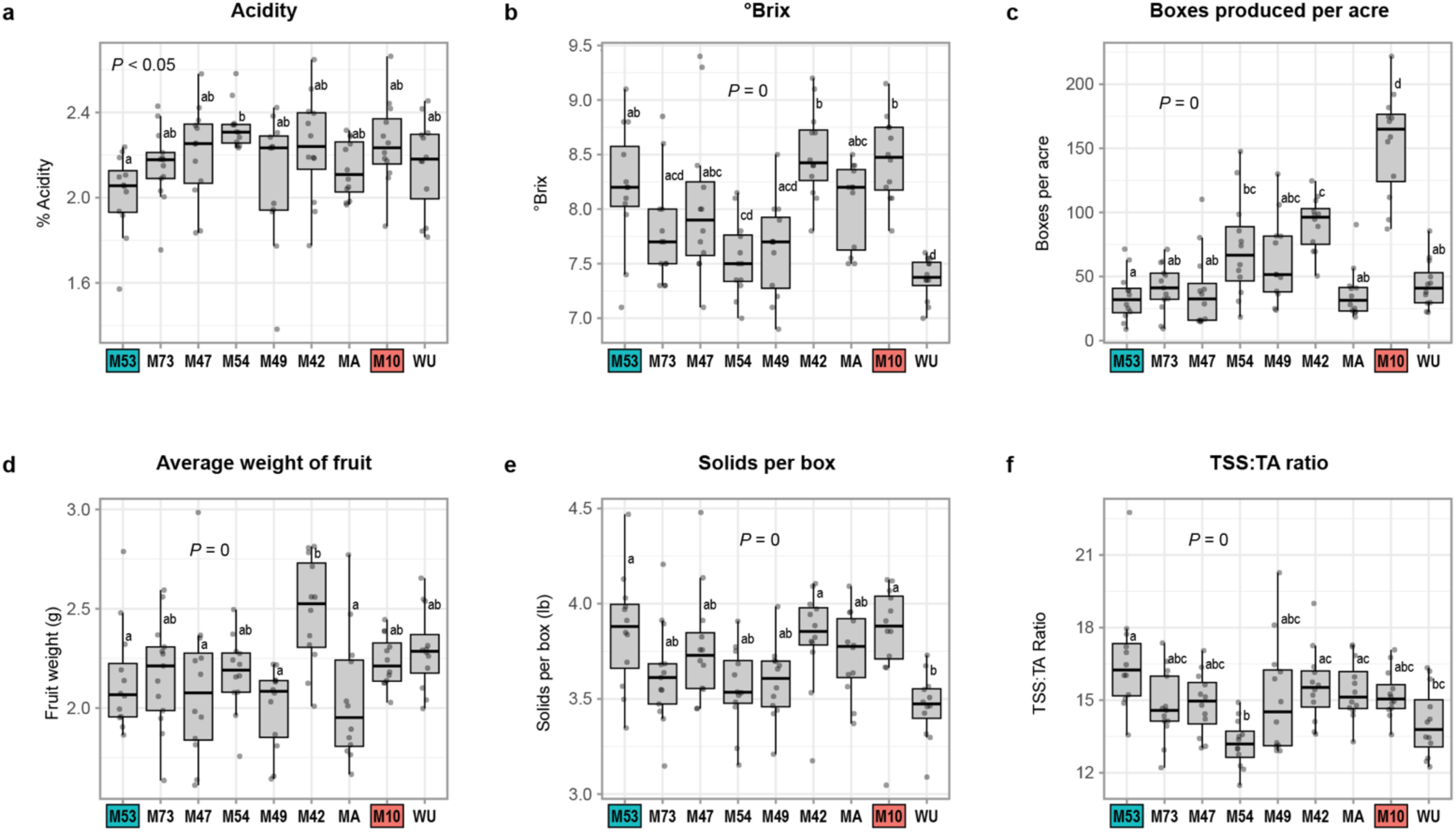
Phase 2 field performance of selected Phase 1 chemistries in Hardee county grove site. Traits include total boxes per acre, mean fruit weight, °Brix, titratable acidity, solids per box, and the TSS:TA sweetness-to-acidity ratio. Treatments M10 and M42 show increases in boxes per acre and average fruit weight, respectively, as compared to M53. *Citrus sinensis* (Hamlin) scions were used, but the grower did not have records for the rootstock. Grower inherited this grove from his father, who passed away.

**Extended Data Fig. 15.**
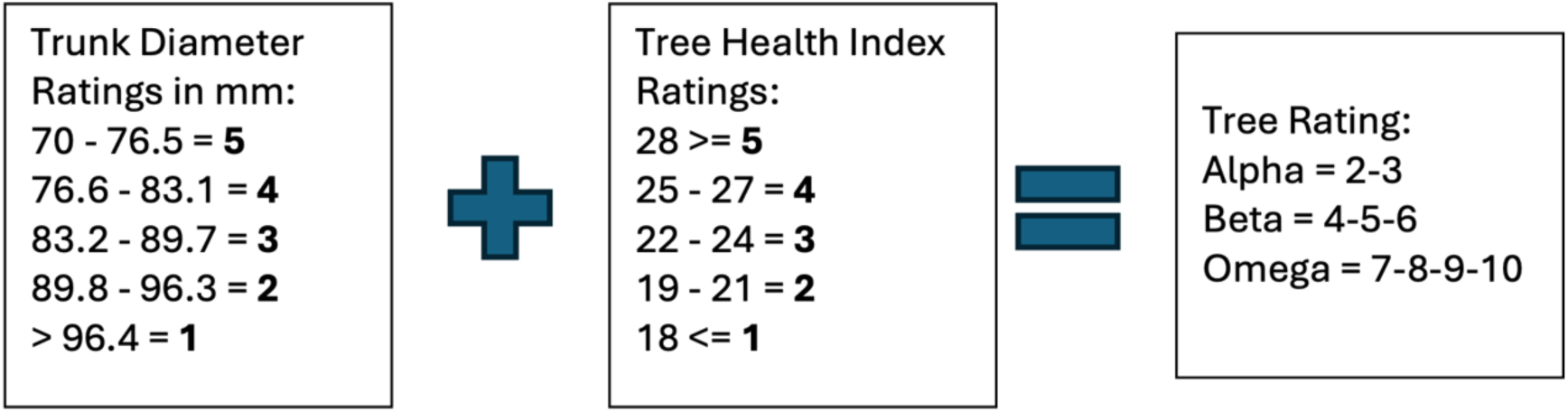
Schematic showing calculations used to classify trees as alpha, beta or omega trees. The group classification was based on the combined score of trunk diameter and tree health index ratings (e.g., if the trunk diameter rating = 1 and tree health index rating = 2, resulting in a total tree rating of 3, then the tree is then classified as Alpha).

## Extended Data Table Titles

**Extended Data Table 1. List of treatments and treatment combinations used in the Phase 1 Grove-First screen.**

**Extended Data Table 2. Pairwise tests against M53 for fruit drop.**

**Extended Data Table 3. Pairwise tests of yield to M53.**

**Extended Data Table 4. Pairwise tests of fruit weight to M53.**

**Extended Data Table 5. Pairwise °Brix comparison against M53.**

**Extended Data Table 6. Pounds solid per box greater than or equal to M53.**

**Extended Data Table 7. Pairwise testing to mean for each tree class with initial rating of alpha, beta or omega in Phase 1 screen.**

**Extended Data Table 8. Power analysis of Phase 1 data. Detectable changes by a one-tailed T-test, given observed variance of each parameter among alpha, beta and omega trees.**

**Extended Data Table 9. Phase 2 Treatment Summary.**

**Extended Data Table 10. Data analysis code headers for Phase 1 analysis.**

## Author Contributions

M.H, and R.P.N. conceived the Grove-First framework, co-supervised the project and wrote the first draft of the paper. W.C.E. provided inspiration for ‘Grove-First’, specific guidance on injection best practices, and guidance on data collection. F.G. coined the term “Grove-First” to describe the framework. N.L., G.L., R.P.N., W.C.E., E.C. and M.H. designed and conducted the experiments. N.L. and G.L. directed the field team. N.L., G.L., E.C., A.M., R.N., M.H., J.H., V.B, L.U., L.R.B., C.E. and M.R. collected the data. M.H., L.J., S.C., R.P.N. and G.L. analyzed the data. M.H., W.C.E., L.J., S.C., G.L., N.L., R.A. and R.P.N. interpretated the data. B.S. developed the “Scully List”, conducted background safety and regulatory research for treatments. M.H. led the treatment team, which consisted of members R.P.N, N.L., G.L, M.P, R.G.S., R.A., C.E., and M.R.. Many co-authors performed trunk injections. E.C. sourced the treatments and prepared treatment solutions for injection. W.C.E., R.A. and R.M. provided access to Phase 2 test sites. B.A. and R.M. contributed treatments to test and guidance on data collection and interpretation. M.H., R.P.N., L.R., M.R., W.C.E.. F.Z., and R.G.S contributed funding. M.R. provided access to the fruit sorting machine and know-how on best practices. L.R. provided guidance on field best practices. L.J. and M.H. conducted the statistical analyses. N.D. conducted the antimicrobial assays and took tree measurements. A.M., G.L., N.L. and E.C. took tree measurements and performed tree health index ratings. All authors read, edited and approved the final the manuscript.

## Supplementary Information

Supplementary Information is available for this paper.

## Correspondence

Correspondence and requests for materials should be addressed to Michelle Heck, Randall P. Niedz or Cody Estes.

## Acknowledgements

We are grateful to Tom Johnson, President of T.J. Biotech, for his generous donation of 400 FlexInject devices used in the study and helpful advice on injection formulations. We thank AgroSource, Inc. personnel for their generous donation of Rectify^TM^ used for OTC-containing injections in the Phase 1 trial and field assistance. We appreciate the expert assistance by Julia Felice (Felice Information Design) for guidance and support with figure conceptualization. We thank Dave Wood (US Horticultural Research Lab) for his assistance with worker safety and chemical disposal associated with the project. We thank Jim Poulos (USDA ARS, Northeast Area Technology Transfer Coordinator, retired) for providing guidance on the best way to transfer the M10 technology to citrus growers. Feedback from our NIFA project advisory board members Stewart Gray (USDA ARS, retired), MaryLou Polek (USDA ARS and California Citrus Research Board, retired), Doug Bournique (Indian River Citrus League), Richard Bennett (California Citrus Research Board and citrus grower, retired), Neil McRoberts (UC Davis), Melinda Klein (California Citrus Research Board), and Rick Dantlzer (Citrus Research and Development Foundation), Board Members of the Indian River Citrus League and members of Florida Citrus Mutual was invaluable to helping develop and implement the Grove-First framework. This project was funded by USDA National Institute of Food and Agriculture (NIFA) Emergency Citrus Research and Development Program, Project # 2020-70029-33176 and USDA Agricultural Research Service projects 6034-21000-020-000-D (Niedz) and 8062-22410-007-00D (Heck). This research was supported in part by an appointment to the Agricultural Research Service (ARS) Research Participation Program administered by the Oak Ridge Institute for Science and Education (ORISE) through an interagency agreement between the U.S. Department of Energy (DOE) and the U.S. Department of Agriculture (USDA). ORISE is managed by ORAU under DOE contract number DE-SC0014664. The Florida Citrus Research and Development Foundation graciously provided funding for grower participation in Phase 2 trials.

